# Pathway-scale variant effect mapping reveals the importance of cell context

**DOI:** 10.64898/2026.10.04.756099

**Authors:** Jessica J. Simon, Sriram Pendyala, A. Felicia Adebanjo, Daniel L. Holmes, Madeline Walsh, Allyssa J. Vandi, Ziyu R. Wang, Melinda K. Wheelock, Jordan L. Opsahl, Aman Shihora, Abbye E. McEwen, Nahum T. Smith, Siobhan O’Brien, Melissa Hopkins, Alan F. Rubin, Alice H. Berger, Lea M. Starita, Dustin J. Maly, Douglas M. Fowler

## Abstract

Variation alters inherent protein properties like thermodynamic stability and catalytic activity, and these intrinsic effects are shaped by the cellular environment. The RAS–MAPK signaling pathway typifies this interplay, with hundreds of proteins acting in concert through regulated states and interactions to transduce signals. To explore how cell context influences variant effects, we measured the abundance and pathway activity of 186,676 variants across 17 MAPK proteins in human cells under diverse conditions, for a total of 503,099 measurements. We discovered 14,412 dominant negative variants, nearly all previously unknown, that disrupt specific domains or interaction interfaces while preserving other functions. MAPK dominant negatives occur at conserved positions and are depleted from the human population relative to other activity-altering variants, representing a distinct constraint on the human genome. Modulating chaperone activity exposed previously hidden effects of genetic variation on protein abundance in human cells, revealing the HSP90 dependence of 67% of missense variants across the pathway. Combining hundreds of experimental and predicted protein interaction structures with variant effect data distinguished functional from non-functional interaction interfaces and revealed how their disruption affected function. We resolved the overlapping KRAS-LZTR1-SOS1 interface into partner-specific and shared binding determinants, identified allosteric networks supporting each interaction, and uncovered distinct mechanisms by which variants promote LZTR1-mediated degradation. Thus, we measure variant effects at pathway scale in human cells, illuminating fundamental MAPK pathway signaling mechanisms and demonstrating that cellular context profoundly shapes the effect of most variants.

## Introduction

The effects of protein-coding variation are shaped by protein structure and intrinsic function as well as the complex cellular context in which proteins fold, localize, and engage interaction partners. The receptor tyrosine kinase (RTK)–RAS–MAPK signaling pathway typifies this interplay, comprising over 30 core and dozens of accessory proteins possessing diverse enzymatic activities and numerous intramolecular and intermolecular interaction sites^1,2^. RTKs receive extracellular growth signals and transduce them by promoting the formation of active RAS-GTP, which is relayed through the RAF–MEK–ERK kinase cascade to drive gene expression programs controlling proliferation and differentiation (**Fig. 1a**). Guanine nucleotide exchange factors (GEFs), GTPase-activating proteins (GAPs), phosphatases, E3 ubiquitin ligases, and scaffolding and adapter proteins link these core components and tune the magnitude and duration of pathway activation. In unstimulated cells, many pathway members are held in an inactive state through intramolecular and intermolecular autoinhibitory interactions. Signal propagation requires the coordinated relief of autoinhibition, spatial reorganization of pathway members, and formation of active signaling complexes. MAPK signaling is required for development in vertebrates, and, in humans, MAPK pathway variants can cause cancer or rare diseases collectively termed RASopathies, illustrating the pathway’s centrality^3–6^. A comprehensive atlas of MAPK pathway variant effects could shed light on the molecular logic of the pathway and reveal how variant effects are shaped by cellular context, but requires measuring variant effects across many pathway members in human cells.

**Figure 1.**
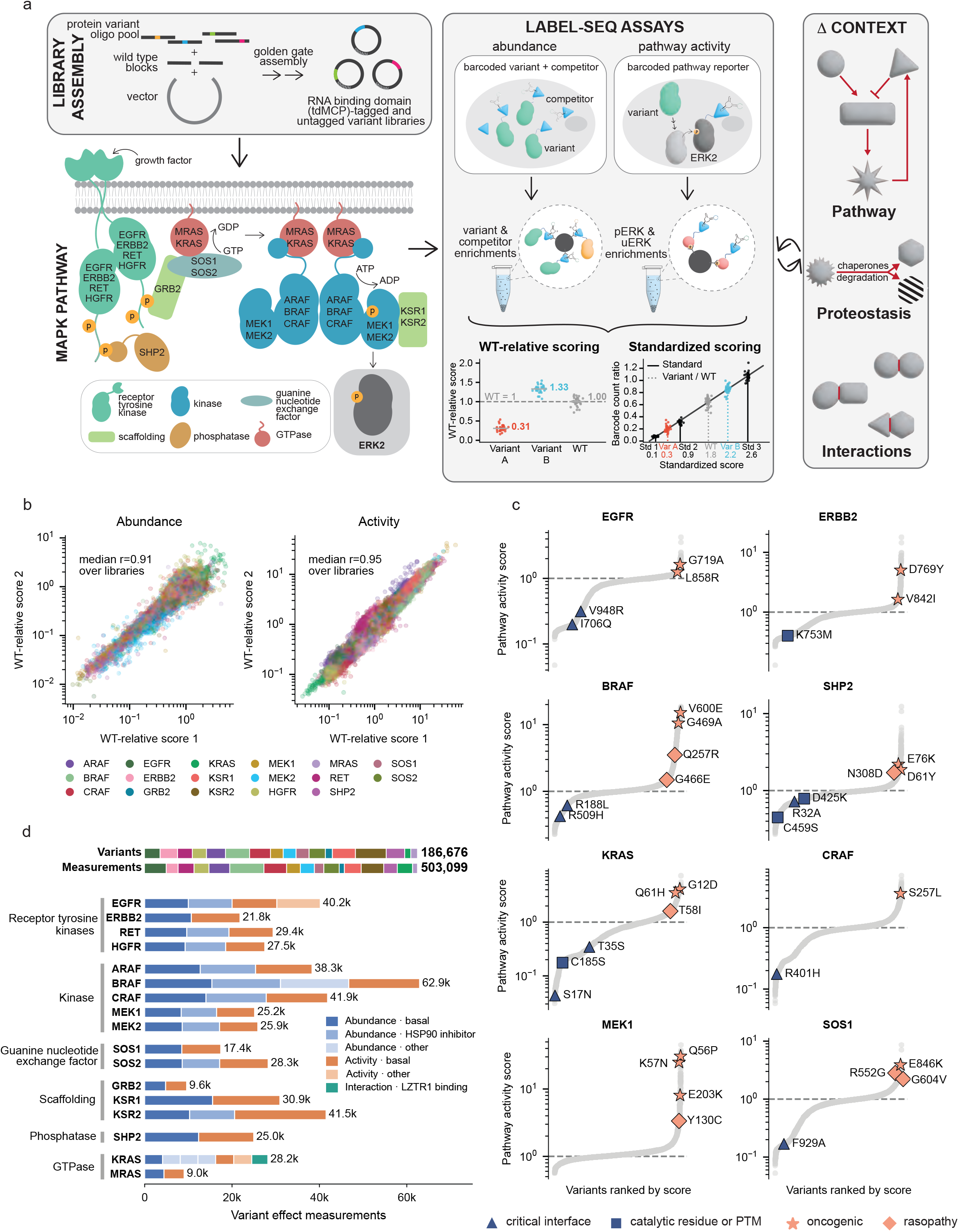
Massively scaled, quantitative profiling of MAPK pathway protein variant abundance and activity. **(a)** Summary of experimental workflow and MAPK pathway. **(b)** Reproducibility of all variant effect scores between two of the three abundance replicates (median over libraries r = 0.91, n = 167,400 variants, left) and pathway activity (median over libraries r = 0.95, n = 183,554, right) assays under basal conditions. Pearson’s r was calculated by linear regression on log_10_-transformed scores. **(c)** Rank plot of pathway activity scores for eight target proteins under basal conditions. Dashed line denotes WT score, marker shape denotes common oncogenic (star)^89^; catalytic position or critical post-translational modification site (square); critical protein–protein interface position (triangle); or RASopathy-associated (diamond) variants. Marker color denotes scores greater than (salmon) or less than (navy) WT. **(d)** Barplot summarising the number of variants per protein quantified in each assay and context. Top bars show total variants or measurements colored by target, as in (b). Bottom bars show measurements per target, colored by assay and context.

To create a MAPK pathway variant effect atlas, we developed synthetic DNA variant cloning (svClone), a scalable variant library synthesis workflow, and combined it with massively scaled LABEL-seq, which converts diverse cellular assays into biochemical enrichments followed by barcode sequencing (**Fig. 1a**)^7^. LABEL-seq leverages the intracellular self-assembly of an RNA-binding domain, tdMCP, with a circular RNA (circRNA) containing a hairpin with high affinity for the tdMCP domain and a short barcode sequence that identifies a co-expressed protein variant. Protein variant properties or functions can be quantified after cell lysis by enrichment of tdMCP-tagged protein fusions, followed by high-throughput sequencing of co-enriched barcodes. Together, svClone and LABEL-seq allowed us to measure the abundance and MAPK pathway activity level for 186,676 synonymous, missense, deletion, and nonsense variants in 17 targeted pathway members. Similarly-scaled variant effect datasets have been collected for small protein domains using cell-free or yeast-based assays that capture folding stability or surrogate functional outputs, which cannot fully recapitulate the pathway context, membrane environments, proteostasis machinery and protein–protein interactions that govern signaling protein function in human cells^8–10^.

Our atlas shows how the biophysical, biochemical, and structural features of variants combine with cellular context to shape the functional consequences of variation across the MAPK pathway. Pathway activity measurements revealed 14,412 (7.7%) variants that suppressed pathway activity below endogenous levels, acting as dominant negatives that are under stronger purifying selection in the human population than either loss- or gain-of-function variants. Abundance measurements for 106,642 variants across 10 targets under HSP90 inhibition reinforced the privileged client status of kinase domains and uncovered the latent dependence of 67% of missense variants across the pathway on HSP90. Combining hundreds of structures of MAPK pathway protein complexes with variant effect data distinguished functional from non-functional interfaces, provided experimental support for 324 predicted interactions, and revealed how disruption of interfaces affected protein abundance and pathway activity. We resolved the overlapping KRAS-LZTR1-SOS1 interfaces into partner-specific and shared binding determinants, identified allosteric networks supporting each interaction, and uncovered distinct mechanisms by which variants promote LZTR1-mediated degradation. Thus, we establish pathway-scale variant effect mapping in human cells, illuminating fundamental MAPK pathway signaling mechanisms and demonstrating that cellular context profoundly shapes the effect of most variants.

## Results

### Mapping variant effects at pathway scale

We selected 17 targets representing a diversity of molecular activities: the receptor tyrosine kinases (RTKs) EGFR, ERBB2, HGFR, RET; the phosphatase SHP2; the adapter GRB2; the guanine nucleotide exchange factors (GEFs) SOS1, SOS2; the GTPases KRAS, MRAS; the RAF kinases ARAF, BRAF, CRAF; the MEK kinases MEK1, MEK2; and the pseudokinase scaffold proteins KSR1, KSR2 (**Fig. 1a, Supplementary Table 1**). These proteins range from 188 to 1,390 amino acids in length, span from membrane-bound to predominantly cytosolic, and contain diverse folded domains as well as disordered regions (**Extended Data Fig. 1a**). Despite this structural and functional diversity, all targets share the ability to control the activation state of ERK, the terminal effector kinase of the MAPK signaling cascade.

We programmed and cloned 202,520 synonymous, missense, deletion, and nonsense variants in the intracellular regions of these 17 targets using synthetic DNA variant cloning (svClone), a method that combines array-synthesized oligonucleotide pools with invariant synthetic DNA blocks to produce full-length open reading frame libraries (**Fig. 1a, Extended Data Fig. 2a**). Each library was cloned into barcoded expression constructs enabling single-copy integration at a genomic landing pad in HEK293 cells, encoding either untagged or tdMCP-tagged variants for use in distinct LABEL-seq assays^7,11^. Across all 44 libraries, 91.1% of sequences corresponded to programmed variants and position-level skewness compared favorably to commercial libraries and other oligonucleotide pool-based methods (**Extended Data Fig. 2b,c,d**)^12^. Thus, svClone enables high-quality site-saturation mutagenesis library cloning on a massive scale.

We used optimized ratiometric LABEL-seq assays to measure two key phenotypes shared by all 17 targets: protein abundance and MAPK pathway activity (**Fig. 1a**). Protein abundance was scored as the ratio of barcoded circRNA enriched by the tdMCP-tagged variant protein to that of a stably expressed tdMCP-tagged competitor protein. We used the phosphorylation state of ERK2, the terminal kinase of the MAPK pathway, as a proxy for pathway activity, scored as the ratio of barcoded circRNA enriched by phosphorylated tdMCP-ERK2 (pERK2) to that enriched by unphosphorylated tdMCP-ERK2 (uERK2) in a sequential pulldown^13^. To enable direct comparison of variant effects across proteins and experiments, every assay included internal abundance or pathway activity standards spanning a broad range of scores. Thus, variant effects were scored on both a wild type (WT)-relative scale, reflecting the variant’s effect relative to the WT sequence of the target protein, and a standardized scale, reflecting the variant’s effect on a common scale across all proteins (**Fig. 1a**).

WT protein abundance and pathway activity spanned two orders of magnitude, while variant abundance and pathway activity spanned three orders of magnitude, underscoring the importance of cross-assay standardization; scores were highly reproducible (median abundance Pearson’s r = 0.91, range 0.57-0.96; median pathway activity Pearson’s r = 0.95, range = 0.66-0.99, **Fig. 1b, Extended Data Fig. 1b,c, Supplementary Table 2**). Pathway activity scores recapitulated the effects of previously characterized oncogenic, RASopathy and reduced activity variants, and single-nucleotide missense variants with increased pathway activity scores were enriched amongst frequently occurring cancer variants (Fisher’s exact test p < 0.01, **Fig. 1c, Extended Data Fig. 1d**). The relative abundance of the WT proteins was consistent with endogenous protein levels in HEK293 cells as determined by quantitative mass spectrometry, supporting the physiological relevance of our system (**Extended Data Fig. 1e**)^14^. In addition to basal abundance and pathway activity measurements, we also evaluated subsets of variants in different cellular contexts, including by inhibiting the HSP90 chaperone, increasing RAS-GTP levels, deleting an E3 ligase adapter protein, and serum starving cells. Thus, combining svClone with LABEL-seq enabled pathway-scale mapping of 503,099 variant effects on protein abundance, MAPK pathway activity and other properties with exceptional dynamic range, data quality, and cross-assay standardization (**Fig. 1d**). These data can be explored at mapk.gs.washington.edu.

### Domain architecture, intramolecular interactions and pathway role shape variant effects on abundance and pathway activity

The effects of the 167,400 profiled variants on abundance differed dramatically across targets and independently of the WT protein abundance (**Fig. 1a, 2a, Extended Data Fig. 3a**). As expected, nonsense variants generally reduced abundance relative to WT for the 11 targets with N-terminal tdMCP tags. However, the proportion of reduced abundance nonsense variants and the magnitude of the abundance decreases they drove depended on the overall domain architecture of the protein and the location of the truncation (**Fig. 2b, Extended Data Fig. 4**). The effects of missense variation on abundance also differed across targets, with MRAS (40.3%) and GRB2 (29.2%) harboring the largest proportions of reduced abundance missense variants and HGFR (8.3%) and MEK1 (8.4%) the smallest. Many missense variants increased abundance, especially in HGFR (15.8%), MRAS (15.4%), and KRAS (9.7%), more than observed for proteins profiled with other multiplexed methods^15–18^. These increased abundance variants were enriched within domains and at protein-protein interaction interfaces (**Extended Data Fig. 3b**), suggesting that many may occur at positions that engage proteins promoting degradation or turnover. Reduced abundance variants were also enriched at buried positions and within domains, while solvent-exposed positions and regions lacking secondary structure harbored a higher proportion of WT-like variants (**Extended Data Fig. 3b**). Proline substitutions were the most detrimental to abundance, and deletions were generally less well tolerated than missense variants, as expected (**Fig. 2a,b, Extended Data Figs. 3c, 4**).

**Figure 2.**
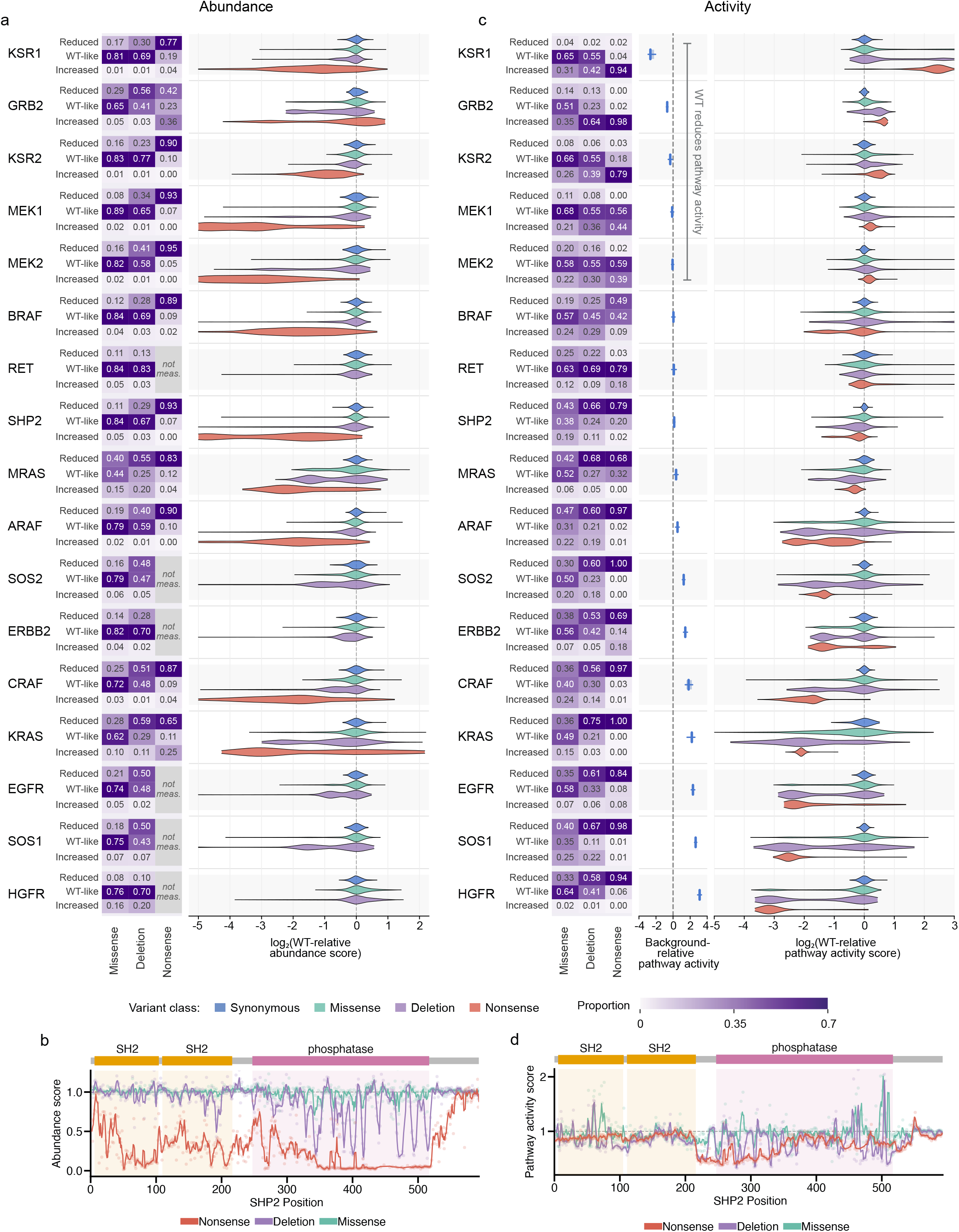
Protein domain architecture, structure, and pathway logic dictate variant effects on pathway activity and abundance. **(a)** Summary of missense, 3-nucleotide deletion, and nonsense variant effects on protein abundance across target proteins under basal conditions, ranked by median synonymous standardized activity score (top, lowest; bottom, highest). Left: heatmap of the proportion of missense, deletion, and nonsense variants per protein classified as reduced, wild type (WT)-like, or increased abundance (see Methods). Right: violin plots of WT-relative abundance scores for synonymous (blue), missense (teal), deletion (purple), and nonsense (red) variants; dashed line denotes WT score. HGFR, RET, ERBB2, SOS2, SOS1 and EGFR lack nonsense abundance scores because they were C-terminally tagged. **(b)** Domain architecture and position mean WT-relative abundance scores for SHP2 variants under basal conditions. Top: colored segments denote domain boundaries. Bottom: line indicates a three position rolling mean; points show individual position means; dashed line denotes WT score. **(c)** WT-relative pathway activity scores plotted as in (a). The box plot (middle) denotes standardized activity scores of synonymous variants only (whiskers, 5th–95th percentile; box, 25th–75th percentile; line, median) normalized to each protein’s empty vector background level (dashed line). **(d)** Pathway activity domain and position plot as in (b).

The effects of the 183,554 profiled variants on pathway activity also differed dramatically across targets (**Fig. 2c**). We measured pathway activity under basal conditions in HEK293 cells where endogenous MAPK pathway activity is low, enabling us to detect increases or decreases in pathway activity. Overall, 26.6% of missense variants reduced pathway activity relative to the WT protein while 19.9% increased activity, with considerable protein-specific variation driven by each target’s domain architecture (**Fig. 2d, Extended Data Fig. 5)**, its role in the pathway, and the effect of exogenous WT expression on endogenous pathway activity. For example, among the 12 targets where the WT protein did not decrease endogenous pathway activity, most nonsense variants reduced pathway activity relative to WT (**Fig. 2c**). RET and BRAF, which minimally affected pathway activity, were exceptions. >20% of SOS and RAF homolog missense variants increased pathway activity, reflecting the extensive autoinhibitory interactions that suppress these targets’ catalytic activity. In contrast, <10% of EGFR, ERBB2, HGFR, and MRAS missense variants increased pathway activity, reflecting their more limited autoinhibitory interactions^19^. Among the five targets whose WT expression reduced pathway activity below endogenous levels, most nonsense variants increased pathway activity relative to WT, consistent with truncated forms of these proteins lacking the capacity to inactivate the pathway. Missense variants of these five targets more often had WT-like pathway activity, consistent with substitutions being less likely than truncations to disrupt their pathway-inhibitory functions.

Across targets, proline substitutions produced the widest range of both reduced and increased pathway activity, highlighting the ability of disruptive substitutions to perturb the functional, pathway-activating state at some locations or the autoinhibited, inactive state at others (**Extended Data Fig. 3d**). Activity-altering missense variants were enriched at buried positions and within domains, and were depleted at solvent-exposed positions and within regions lacking secondary structure, consistent with disruption of either pathway-activating or autoinhibited states (**Extended Data Fig. 3e**).

Integrating abundance and pathway activity measurements revealed that, as expected, variants with reduced pathway activity also generally had reduced abundance (**Fig. 3a**). However, variants that increased pathway activity also generally had reduced abundance, suggesting that alterations in activity in either direction arise predominantly from disruption of stabilizing interactions. Reduced-activity variants were enriched at buried positions, consistent with disruption of interactions that stabilize the pathway-activating state whereas increased-activity variants were enriched at interdomain contacts, consistent with disruption of autoinhibitory interactions (**Fig. 3b**). For example, at buried positions in the CRAF kinase domain C-lobe (region 2; **Fig. 3c,d**), the majority of variants reduced activity and abundance, suggesting inactivation through disruption of the pathway-activating state. In contrast, variants at positions that participate in two major networks of autoinhibitory interactions (regions 1 and 3; **Fig. 3c,d**) predominantly increased activity and reduced abundance^20^. Consistent with autoinhibition providing stabilization, transient upstream pathway activation, which disrupts BRAF’s autoinhibited state, reduced the abundance of WT BRAF and most missense variants relative to unstimulated cells (**Extended Data Fig. 6a,b**). Thus, the effects of variants on MAPK pathway protein abundance and activity are tightly linked and dependent upon the domain architecture of each protein, its intramolecular interactions, its role in the pathway, and the activation state of the pathway within the cell.

**Figure 3.**
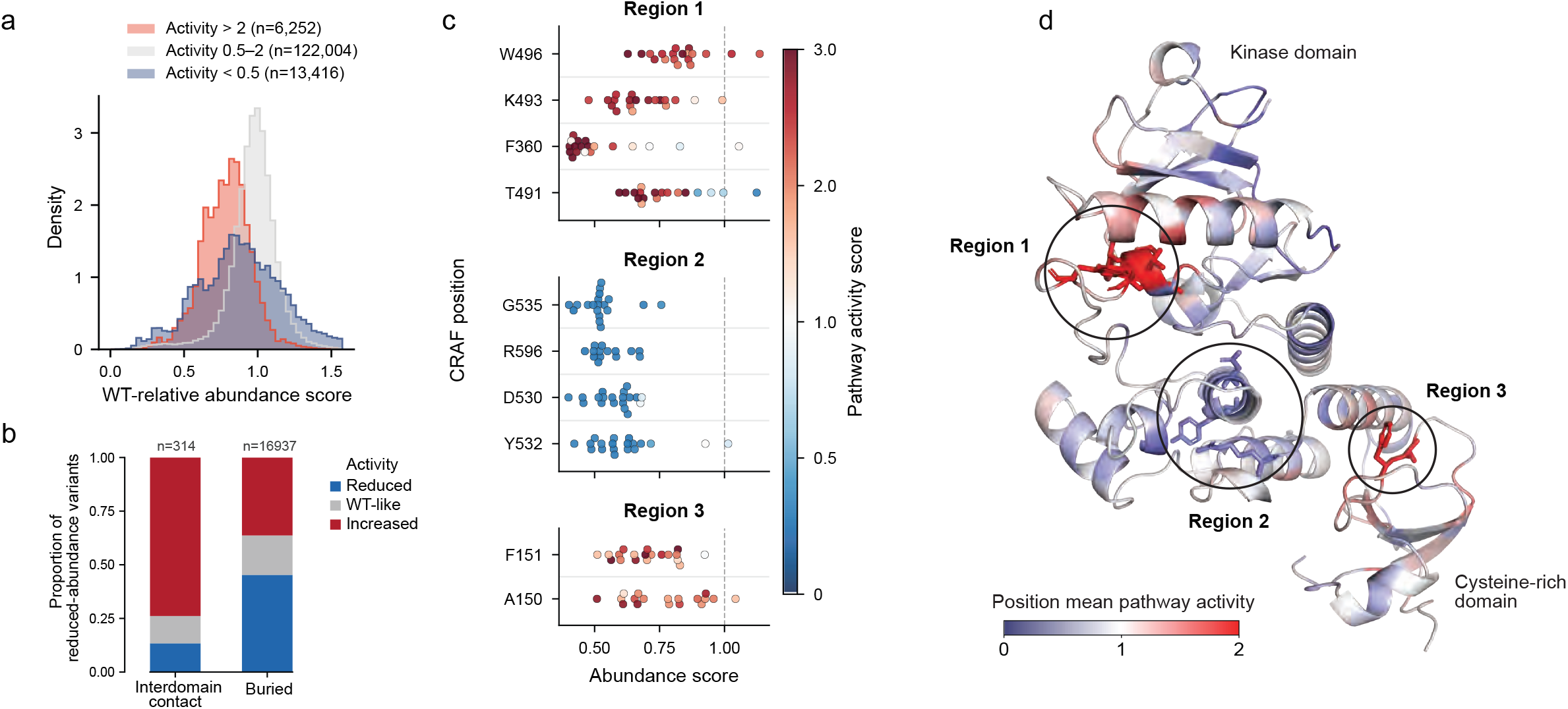
Activity-altering variants disrupt stabilizing interactions and generally reduce abundance. **(a)** Distribution of WT-relative missense abundance scores under basal conditions, stratified by activity score. **(b)** Proportion of reduced abundance missense variants with reduced, WT-like, or increased activity, stratified by structural context (see Methods for classifications). **(c)** WT-relative abundance scores for missense variants in three regions of CRAF under basal conditions, colored according to WT-relative pathway activity score; dashed line denotes WT score. **(d)** CRAF structure (PDB 9MMP) showing the kinase and cysteine-rich domains, colored by position mean pathway activity score. Region 1 = autoinhibitory turn within the activation loop, region 2 = buried positions within the kinase domain C-lobe, region 3 = interface between the cysteine-rich domain and kinase domain C-lobe.

### Dominant negative variants are pervasive in the MAP kinase pathway and depleted from the human population

Variants can increase or decrease overall protein function, leading to gain or loss of pathway activity. However, variants can also disrupt some aspects of protein function while leaving others intact, acting as dominant negatives (DNs) that reduce pathway activity below endogenous levels by interfering with the activity of WT protein^21^. For example, KRAS S17N traps the protein in an inactive state while preserving upstream GEF interactions, thereby suppressing endogenous pathway activity^22,23^. Only a handful of other DN variants in ARAF, EGFR, ERBB2, SHP2 and KRAS have been identified, variant effect predictors generally underperform on DN variants, and systematic mapping of DN variants has been confined to single proteins or short peptides rather than conducted for full-length proteins at pathway scale^24–32^.

Our standardized pathway activity measurements enabled us to identify biochemical DN variants that, when expressed in the context of the genomic copies of the target protein, suppressed pathway activity below endogenous levels (**Fig. 4a**). Among the 12 WT proteins that did not decrease pathway activity, we identified 14,412 such DN variants: 11,514 missense, 1,592 deletion, and 1,306 nonsense (**Fig. 4b**). Missense and single-position deletion DN variants were enriched in domains (odds ratio, OR, of 3.7 and 4.5, Fisher’s exact test FDR-corrected q < 10^-300^ and 1.3 × 10^-61^, respectively) and were depleted from solvent-exposed positions with no annotated function (q < 10^-300^ and 6.2 × 10^-75^; **Fig. 4c**, **Extended Data Fig. 7a**). They were also enriched at active sites (ORs of 3.1 and 2.3, q = 2.0 × 10^-277^ and 4.2 × 10^-12^) and at protein-protein interaction interfaces (ORs of 2.3 and 2.3, q < 10^-300^ and 2.8 × 10^-23^), as previously shown for DNs broadly (**Fig. 4c,d**, **Extended Data Fig. 7a**)^33–35^. Lastly, they were enriched at buried, non-active site positions (ORs of 2.1 and 2.5, q = 4.4 × 10^-291^ and 1.1 × 10^-36^). Across all 12 targets, 82% of missense or single-position deletion DN variants were at active site, buried or interface positions, with some positions occupying more than one category (**Fig. 4d**). In contrast, truncating DN variants were not significantly enriched at active sites (OR 1.01), buried positions (OR 0.98) or interfaces (OR 1.06), instead creating protein fragments that likely interfere with the same activity, folding and interaction functions by removing or disrupting key domains (**Fig. 4c**). Thus, by measuring pathway, rather than protein, activity, we discovered a massive number of MAPK pathway DN variants which were previously unknown despite decades of intensive study including numerous mutational scans of pathway members^36–52^. DN variant locations suggested their mechanism, and their distribution differed considerably between proteins.

**Figure 4.**
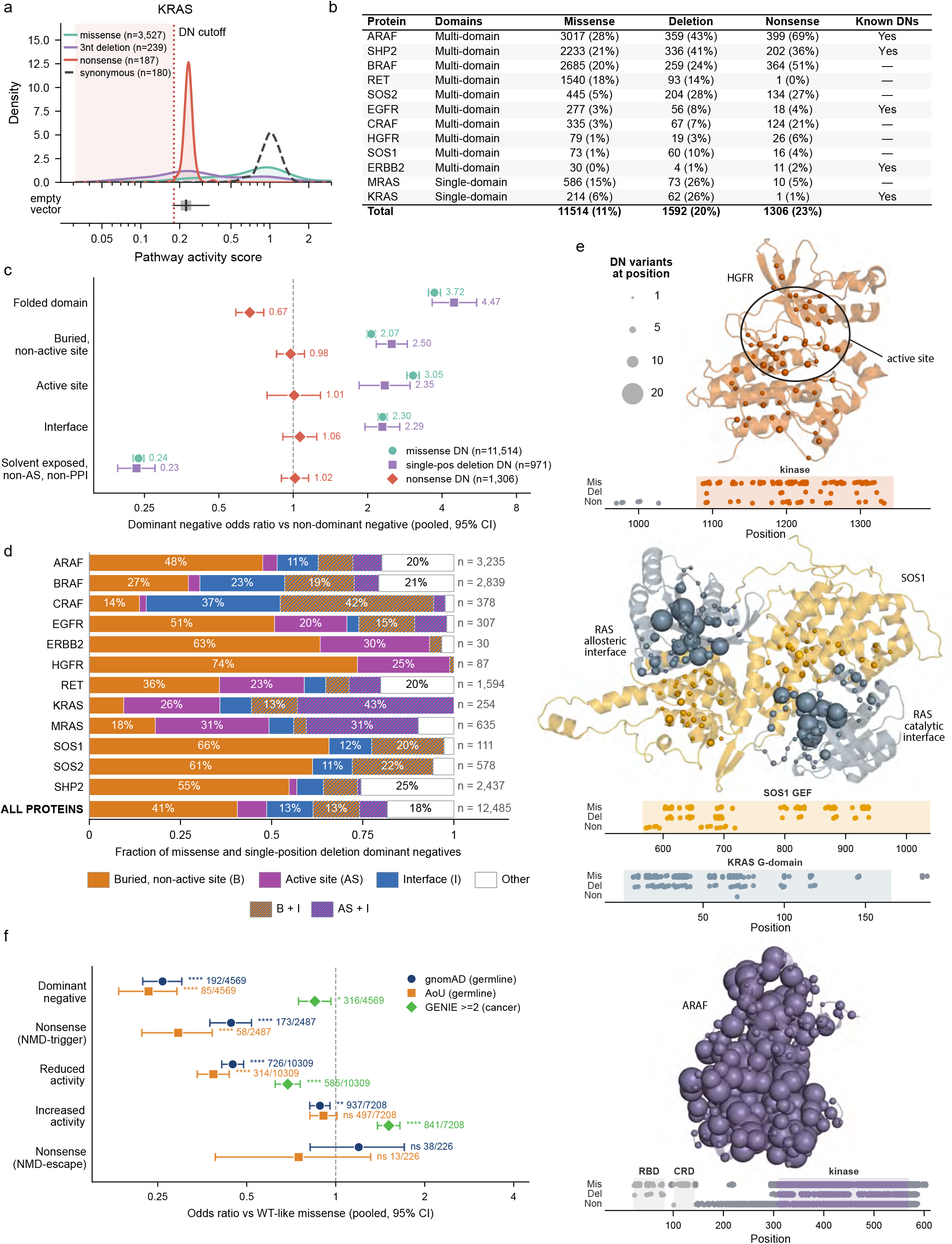
∼14,000 variants across the MAPK pathway act in a dominant negative fashion and are depleted from the human population. **(a)** Top: pathway activity score distributions for KRAS synonymous, missense, 3-nucleotide deletion, and nonsense variants. Bottom: box plot of bootstrapped mean pathway activity scores from 500 samples, each with 10 empty vector barcodes; whiskers denote 2.5th to 97.5th percentiles, box spans 25th to 75th percentiles, vertical black line denotes median of bootstrapped means. Vertical red dotted line denotes the 2.5th percentile of bootstrapped means, used as the dominant negative threshold. **(b)** Table of dominant negative (DN) variants per target annotated by variant type. **(c)** Enrichment or depletion of missense, single-position deletion, and nonsense DN variants in different protein structure categories relative to non-DN variants; bars denote the 95% confidence interval. **(d)** Fraction of missense and single-position deletion DN variants in each target categorized as buried, non-active site (B), active site (AS), interface (I), in multiple categories or in none of these categories (other, see Methods for detail). **(e)** DN variants shown on HGFR kinase domain (top, PDB 3DKC), HRAS-SOS1-HRAS complex (middle, PDB 1NVU; gray = HRAS with spheres showing KRAS DNs at analogous positions, wheat = SOS1), or ARAF kinase domain structures (bottom, PDB 9AXM). Sphere size indicates number of missense and single-position deletion variants at each position, positions with at least one missense (Mis), single-position deletion (Del), and nonsense (Non) DN variants are shown in the dot plot below each structure. **(f)** Enrichment or depletion of different types of single-nucleotide missense or nonsense variants in the gnomAD, AllOfUs or GENIE databases, relative to variants with WT-like pathway activity, with counts shown. Bars denote the 95% confidence interval; asterisks denote significance by two-sided FDR-corrected Fisher’s exact test (**** q < 0.0001, *** q < 0.001, ** q < 0.01, * q < 0.05, ns, not significant).

For example, in HGFR, missense and single-position deletion DN variants were enriched in the kinase active site (OR of 2.8, q = 1.5 × 10⁻^4^; **Fig. 4d,e**, **Extended Data Fig. 7a**). HGFR DN variants that disrupt kinase activity likely form heterodimers with endogenous WT HGFR but are incapable of participating in activating trans-autophosphorylation. For SOS1 and KRAS, DN variants were enriched on both sides of the catalytic interface (OR of 4.2 for KRAS, 5.2 for SOS1, q = 6.1 × 10^⁻23^, 2.8 × 10^-10^), and in CRAF, DN variants were enriched at the dimerization interface (OR of 15.0, q = 1.8 x 10^-118^; **Fig. 4d,e**, **Extended Data Fig. 7a,b**). Variants that disrupt these interfaces likely prevent SOS1 from promoting the formation of active GTP-bound RAS and CRAF from forming active dimers, while inhibiting endogenous pathway activity by sequestering signaling partners through their intact domains. Buried SOS1 DN variants also form a network connecting the allosteric SOS1 interface to the catalytic interface (**Extended Data Fig. 7c**), suggesting that disruption of allosteric communication represents an additional mechanism of dominant negativity^53^. In ARAF, missense and deletion DN variants were enriched at buried sites, 95% of them in the kinase domain (OR of 4.1, q = 6.3 × 10^-236^; **Fig. 4d,e**). Most ARAF nonsense DN variants were also in the kinase domain, and all but one spared the RAS-binding and cysteine-rich domains. ARAF DN variants may therefore act by disrupting activating interactions and catalytic activity through the unfolding of the kinase domain, without altering RAS binding. Thus, DN variants can act by disrupting active sites, domains or interaction interfaces while leaving other functions intact, thereby inhibiting endogenous pathway activity.

These biochemical DN variants inhibit pathway activity when expressed in HEK293 cells, while total loss of MAPK pathway activity results in embryonic lethality in non-human vertebrates^5,54^. Therefore, we hypothesized that our DN variants would be depleted in the human population. Indeed, missense and single-position deletion DNs occur at positions that are more evolutionarily constrained than other variants (pooled Mann-Whitney FDR-corrected q < 5.9 × 10^-81^; **Extended Data Fig. 7d**). Moreover, single-nucleotide missense DN variants are significantly depleted in the genome aggregation database (gnomAD) compared to variants that did not alter MAPK pathway activity (pooled OR 0.26, Fisher’s exact test FDR-corrected q = 4.1 × 10^-98^; **Fig. 4f**, **Extended Data Fig. 7e**)^55^. This depletion replicated in the smaller All of Us (AoU) database (pooled OR 0.23, q = 2.3 × 10^-57^)^56,57^. DN variants are also more depleted than increased pathway activity variants (pooled OR 0.29 in gnomAD, 0.26 in AoU; q = 3.2 × 10^-61^ and 3.6 × 10^-38^) or reduced pathway activity variants (pooled OR 0.58 and 0.61, q = 1.8 × 10^-11^ and 6.8 × 10^-5^), and, in gnomAD, than nonsense variants that trigger nonsense-mediated decay and thus result in total loss of function (pooled OR 0.59 and 0.79, q = 1.7 × 10^-6^ and 0.21; **Extended Data Fig. 7e**). Lastly, DN variants are not enriched in human cancers in the AACR GENIE database, suggesting that they generally do not paradoxically activate the MAPK pathway like kinase-impaired BRAF type III variants (pooled OR 0.85; **Fig. 4f**, **Extended Data Fig. 7e**)^58,59^. Thus, DN variants in the MAPK pathway are pervasive, occurring at conserved sites and appearing to be under strong purifying selection in the human germline, consistent with the importance of MAPK signaling for development. Our results suggest that DN variants are far more common than previously appreciated and may play a major role in shaping human genetic variation.

### Chaperone activity shapes the abundance of most missense variants

Variant effects on protein abundance are a product of intrinsic effects on the stability of the folded state and the action of cellular proteostasis machinery which can stabilize, mature, or degrade proteins. In 14 of our 17 targets, 25% or fewer missense variants reduced abundance, compared to 25-40% for other non-MAPK proteins (**Fig. 2a**)^15–17,60–62^. Many MAPK pathway proteins interact with HSP90, which we hypothesized could be responsible for this mutational tolerance^63–65^. To test this hypothesis, we selected 10 of the 14 tolerant targets spanning a range of HSP90 interaction strengths, and repeated abundance assays in the presence of pimitespib, an HSP90 inhibitor (HSP90i; **Fig. 5a**)^66^. Seven of 10 WT proteins decreased in abundance upon HSP90 inhibition, identifying them as HSP90 dependent. The magnitude of their dependence was consistent with their previously determined HSP90 interaction status (**Fig. 5b**, **Extended Data Fig. 8a**)^63,65^. 67% of 91,584 missense variants across targets were also dependent upon HSP90 compared to only 10.0% of 2,656 nonsense variants, consistent with HSP90 primarily rescuing the folding of destabilized, full-length proteins (**Fig. 5b**)^67,68^. Most variants of highly HSP90 dependent proteins were themselves dependent, whereas variants of moderately or lowly dependent proteins had a smaller proportion of dependent variants (**Extended Data Fig. 8a**).

**Figure 5.**
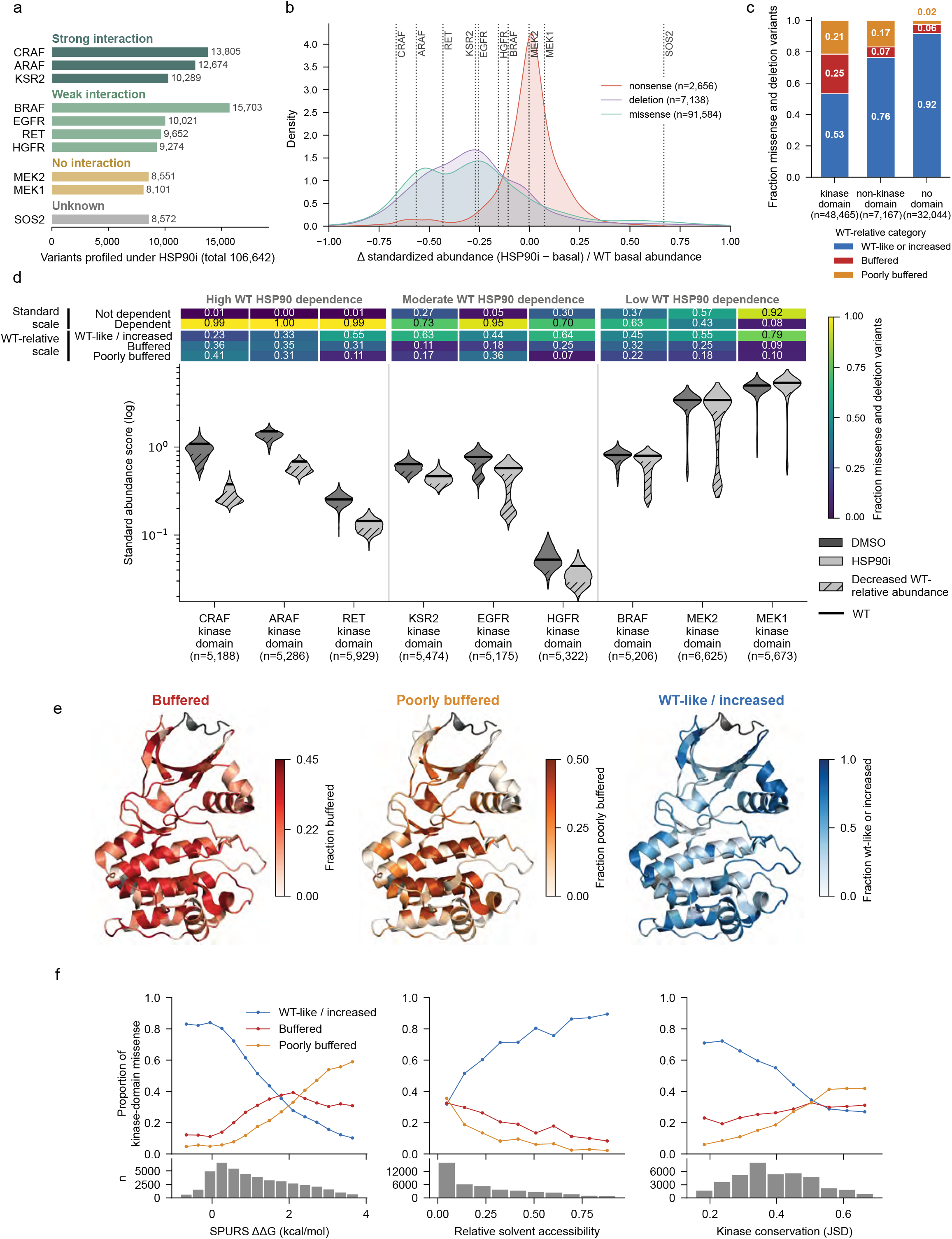
HSP90 shapes the function of many MAPK pathway proteins and their variants, especially in kinase domains. **(a)** Barplot summarizing variant abundance measurements under HSP90 inhibition (HSP90i), across 10 targets, grouped by reported HSP90 interaction status^65^. **(b)** Distributions of missense, 3-nucleotide deletion and nonsense variant change in abundance under HSP90 inhibition, normalized to the wild type (WT) protein abundance under basal conditions for all profiled targets. Dotted lines denote WT target change in abundance. Only variants with both basal and HSP90-inhibited abundance measurements are shown. **(c)** Proportion of missense and deletion variants in each HSP90 WT-relative buffering class in kinase domains, non-kinase domains or no domain across the 9 kinase domain-bearing targets profiled (SOS2 excluded). **(d)** Summary of basal and HSP90i kinase domain missense and deletion abundance scores. Top: heatmap showing the fraction of variants that are dependent or not dependent on HSP90 (standardized scale) and buffered, poorly buffered, or WT-like/increased (WT-relative scale, see Methods for classifications). Colors indicate the fraction of variants in each category. Bottom: violin plots showing abundance score distributions for each target, grouped by WT target dependence. Bars indicate the WT score, violins are trimmed to the 0.25th–99.75th percentile for display. Hatching indicates the portion of the variant distribution that is low abundance (see Methods for classification). **(e)** Mean fraction of buffered (left), poorly-buffered (middle) or WT-like/increased (right) missense variants at each aligned position across the nine profiled kinase domains, projected onto the kinase domain of Aurora-A (PDB 3E5A)^90^. Shade indicates fraction. Aurora-A positions not aligned to other kinases are gray. **(f)** Top: proportion of kinase domain missense variants in each buffering class, binned by SPURS-predicted variant thermodynamic stability relative to WT (left), relative solvent accessibility (middle), and kinase-paralog sequence conservation (right, computed using Jensen–Shannon divergence on the human kinome alignment)^90,91^, pooled across the nine targets. Bottom: variant counts in each bin.

To disentangle the HSP90 dependence of each variant from the dependence of its WT protein, we classified each variant’s abundance relative to synonymous variant abundance measured in the same condition, thereby estimating the degree of chaperone buffering for each variant. Missense variants fell into three categories: variants which had WT-like or increased abundance regardless of HSP90 activity; HSP90-buffered variants, which had WT-like or increased abundance under basal conditions but had reduced abundance relative to WT under HSP90 inhibition; and poorly buffered variants, which had reduced abundance relative to WT regardless of HSP90 activity (**Extended Data Fig. 8a**). Overall, 15.7% of missense variants were buffered by HSP90 and 12.6% were poorly buffered. Within kinase domains, 24.4% of missense variants were buffered compared to 6.8% in non-kinase domains or outside domains, reflecting the preferential rescue of destabilizing variants in kinase domains by HSP90 (**Fig. 5c**). The degree of kinase domain variant buffering was related to the WT protein’s HSP90 dependence (**Fig. 5d**). For example, WT CRAF, ARAF and RET were highly dependent on HSP90, and 33.8% of kinase domain missense variants of these proteins were buffered whereas KSR2, EGFR, HGFR, BRAF, MEK2 and MEK1 had moderate or low dependence on HSP90, and 19.8% of kinase domain variants were buffered. However, while MEK1 and MEK2 share 87% kinase domain sequence identity and similar dependence on HSP90, they had large differences in the fraction of dependent (7.8% vs 42.2%) and buffered (8.8% vs 24.8%) kinase domain missense variants. Surprisingly, SOS2, along with 74.9% of SOS2 missense variants, increased in abundance upon HSP90 inhibition, suggesting that HSP90 or an HSP90-dependent protein mediates degradation of SOS2 (**Fig. 5b**, **Extended Data Fig. 8a**). Thus, MAPK pathway variants display diverse patterns of buffering by HSP90 dominated by the domain in which they occur and the dependence of their WT protein. Without HSP90 activity, the proportion of missense variants that reduced abundance relative to WT rose from 15.8% to 28.3% (21.9% to 43.4% in kinase domains), including among targets whose WT sequence was not HSP90-dependent, indicating that mutational tolerance is conferred widely by HSP90 activity.

To shed light on why some kinase domain variants were buffered and others were not, we analyzed the correlation of 49 biophysical, structural, and biological factors with buffering (**Extended Data Fig. 8b**). The predicted change in variant thermodynamic stability, along with sidechain burial and conservation of the mutated position, were the most strongly correlated with buffering, suggesting that destabilization of folded domains drives buffering (**Fig. 5e,f**)^69^. Variants predicted to cause large decreases in thermodynamic stability were less likely to be buffered than variants that caused modest decreases in predicted stability, extending previous observations that HSP90 preferentially buffers partially destabilized variants, especially in kinases where activation is destabilizing^70,71^. Thus, variants in kinase domains are privileged HSP90 clients, and buffering of non-kinase domain variants may be due to exposure or destabilization of kinase domains. Beyond kinase domains, general instability, rather than specific sequence or structural motifs appears to be a strong driver of client status. Across ∼100,000 variants of 10 proteins, we demonstrate that, in addition to HSP90 interaction, functional dependence on HSP90, in this case for abundance, is widespread^63,65,72^.

### Systematic mapping of MAPK pathway interactions using variant effect data

In addition to interacting with HSP90, MAPK pathway proteins are governed by interactions controlling autoinhibition, signaling activity, feedback, localization, and scaffolding, with ∼2,500 interactions involving ∼1,800 partner proteins reported for our 17 targets^73,74^. Structures of these interactions are rapidly accumulating, with 343 experimentally determined structures in the Protein Data Bank (PDB) and 1,126 predicted structures from proteome-wide *in silico* screens containing interfaces for 575 unique interactions involving our targets (**Fig. 6a**)^75,76^. These structures suggest how interactions may modulate function, but they can be wrong, capture only static snapshots, and cannot establish which interface positions are functionally important or whether an interaction modulates protein function^77^. We hypothesized that combining structures with variant effect data collected in a human cell context could identify functional interfaces and reveal how disrupting each interface position alters protein abundance and pathway activity.

**Figure 6.**
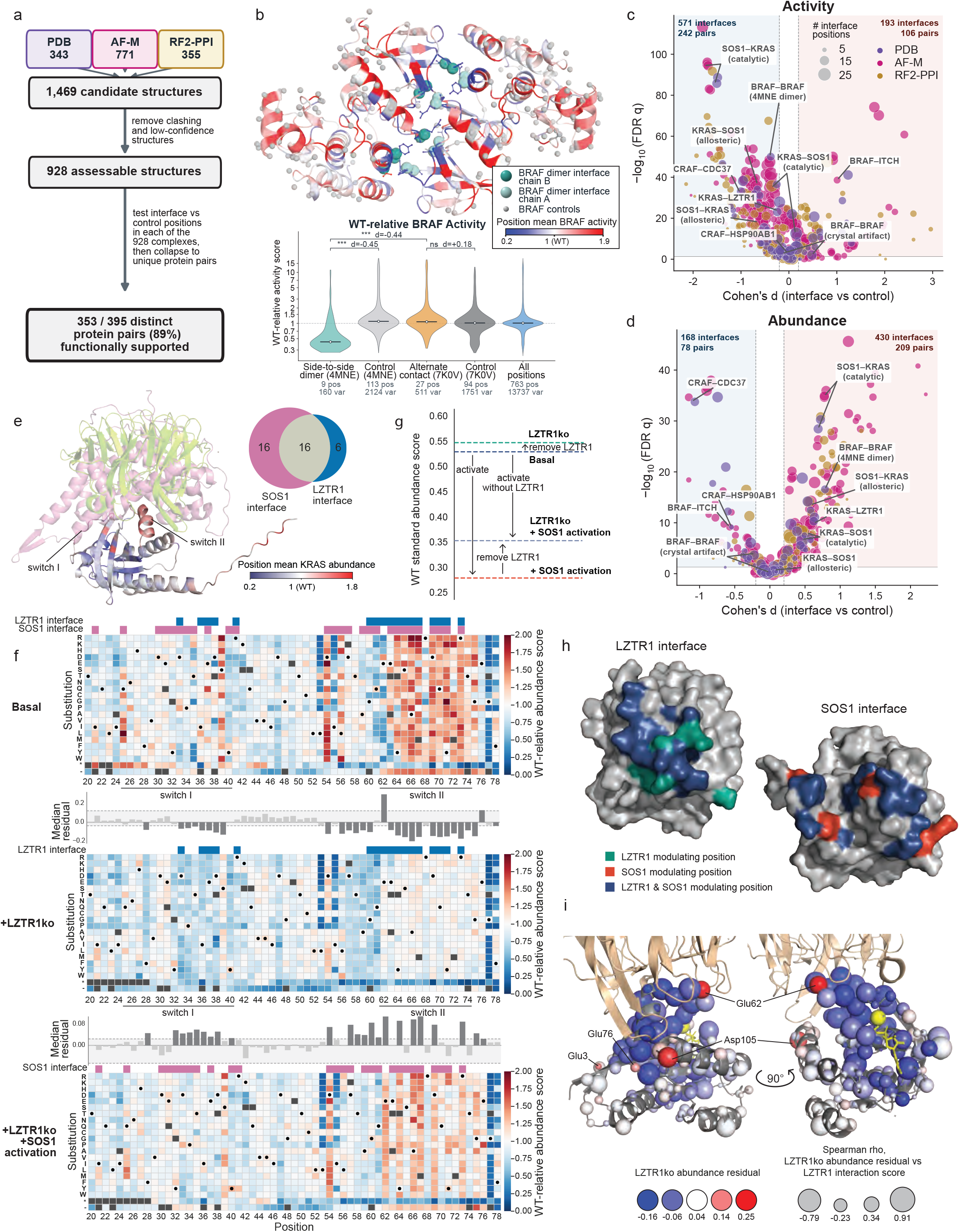
Massive-scale variant effect data reveal the functional importance of protein-protein interaction interfaces. **(a)** Workflow for intersecting variant effect data and experimental or predicted complex structures (AF-M=AlphaFold-multimer, RF2-PPI=RoseTTAFold2-PPI). 353 of 395 distinct pairs of proteins had at least one structure that was functionally supported (q < 0.05 and |Cohen’s d| > 0.2, see Methods). **(b)** Top: active BRAF dimer structure (4MNE) colored by position mean WT-relative pathway activity score. Spheres denote dimer interface (teal, shaded by chain) and control positions (gray). Bottom: WT-relative activity score distributions for the 4MNE (dimer) and 7K0V (crystallographic) interfaces, controls, and all positions. Brackets indicate statistical comparisons (*** indicates Mann-Whitney U-test FDR-corrected q < 0.001), Cohen’s d value shown for comparison. The number of variants (var) and positions (pos) are indicated below each group. **(c)** Volcano plot of Cohen’s d (x-axis) against FDR-corrected q-value (y-axis) across the 1,088 scored interfaces (one per profiled protein in the 928 assessable complexes), colored by the source of the structure. Complexes of the same protein pair and source with overlapping interfaces (intersection-over-union ≥ 0.3) are merged into one point at their median Cohen’s d and geometric mean p-value, then FDR correction is applied across the merged points. Point size denotes the number of interface positions. **(d)** Volcano plot as in (c), but for abundance scores. **(e)** Bottom left: position-mean abundance scores mapped onto the structure of KRAS. Top left: overlay of SOS1 (pink, PDB 1NVU) and LZTR1 (green, 9MF0) bound to HRAS and KRAS, respectively. Right: Venn diagram showing the number of RAS interface positions within 4 Å of SOS1 (magenta), LZTR1 (blue), or both (gray), based on the co-crystal structures. **(f)** Variant effect maps showing KRAS WT-relative abundance scores at positions 20–78 under three conditions: basal, LZTR1-knockout (LZTR1ko), and SOS1 activation in LZTR1ko cells. Dots indicate the WT amino acid; dark gray indicates missing data; (-) indicates single-position deletions; (*) indicates truncations. LZTR1 (blue) and SOS1 (pink) interface positions from (e) are shown above each map. Bar plots represent median per-position residuals from linear regressions of LZTR1ko versus basal variant abundance scores (middle) or SOS1 activation in LZTR1ko cells versus LZTR1ko variant abundance scores (bottom). The shaded gray region represents the 5-95th percentile range of synonymous variant residuals. Positions with median residuals outside this range, denoted by dark gray bars, are defined as LZTR1- and SOS1-modulating. See Extended Data Fig. 10d for residual plots. **(g)** Standardized abundance score of WT KRAS across four conditions: basal (navy), LZTR1ko (teal), SOS1 activation (red), and SOS1 activation in LZTR1ko cells (lavender). **(h)** Interface positions at which KRAS variants modulate the interaction with LZTR1 only (green), SOS1 only (red) or both (blue), projected onto structures of KRAS bound to LZTR1 (left, PDB 9MF0) and of HRAS bound to SOS1 (right, PDB 1NVU); LZTR1 and SOS1 are not shown, for clarity. **(i)** KRAS bound to LZTR1 (PDB 9MF0), with sphere color representing the median missense residual at each position from the regression of LZTR1ko abundance scores versus basal abundance scores. Sphere size denotes the Spearman correlation between the LZTR1ko abundance residual and LZTR1 interaction score.

To test this hypothesis, we first removed structures with steric conflicts or whose interfaces occurred within regions we did not mutagenize, yielding 928 structures comprising 395 unique protein pairs (**Extended Data Fig. 9a**). For each structure, we compared the effects of variants at interface positions to a control set of variants at proximal, solvent-exposed positions, enabling us to distinguish functional and non-functional interfaces (**Fig. 6a**, **Extended Data Fig. 9b,** see Methods). For example, variants in the well-studied BRAF activating dimer interface (4MNE) reduced pathway activity and increased abundance (Cohen’s d of −0.45 and 0.90, Mann-Whitney U-test FDR-corrected q = 2.2 × 10^-55^ and 4.1 × 10^-18^), whereas variants in an artifactual crystallographic interface from a separate structure (7K0V) had negligible effects on function (Cohen’s d of 0.18 and −0.14; **Fig. 6b**, **Extended Data Fig. 9c,d**). Across all complexes, 328 (83%) protein pairs had interface variants with larger effects on pathway activity than controls, including 44 of 53 pairs with experimental structures (**Fig. 6c, Extended Data Fig. 9b**), whereas 275 (70%) protein pairs had interface variants with larger effects on abundance than controls, including 36 of 53 pairs with experimental structures (**Fig. 6d**).

Interface variants often had opposing effects, reducing pathway activity at 75% of functional interfaces and increasing protein abundance at 72% of functional interfaces (**Fig. 6c,d**). This pattern is consistent with interface variants preventing assembly of active signaling complexes, reducing pathway activity while stabilizing the target protein in its autoinhibited state and increasing abundance, as exemplified by the BRAF dimer interface (**Fig. 6b**). Interface variants also revealed direct interaction-mediated modulation of activity and abundance. Two examples are CRAF’s interfaces with CDC37 and HSP90, where variants reduced CRAF abundance (Cohen’s d of −0.67 and −0.84, q = 1.7 × 10^-18^ and 3.7 × 10^-18^, respectively), consistent with CRAF’s chaperone dependence (**Figs. 5b, 6d**, **Extended Data Fig. 9e,f**)^78,79^. A speculative example is BRAF’s predicted interface with the E3 ubiquitin ligase ITCH, where variants reduced abundance and markedly increased pathway activity (Cohen’s d of −0.60 and 1.08, q = 1.1 × 10^-10^ and 2.2 × 10^-41^), providing a plausible structural basis for ITCH’s regulation of BRAF (**Fig. 6c,d**, **Extended Data Fig. 9g,h**)^80^. Overall, 307 of the 353 protein pairs with functional interfaces lacked an experimental structure, 187 were not in curated interaction databases^73,74^, and 83 engaged a partner with a verified drug or chemical probe^81^, highlighting the utility of using variant effect data to dissect predicted interactions (**Supplementary Table 3**). These complexes can be further explored at mapk.gs.washington.edu.

However, unlike the BRAF-ITCH, CRAF-CDC37, and CRAF-HSP90 interfaces, 79% of the interfaces we analyzed overlapped with other interfaces, complicating the attribution of a variant’s effect at the shared interface to a single interaction (**Extended Data Fig. 9i**). For example, the 98 interaction partners of KRAS form 22 overlapping, functionally related groups of interfaces (**Extended Data Fig. 9i,j**). The KRAS switch II helix (positions 64-75) harbors the most promiscuous binding surface among all 17 targets, with 56 KRAS partners predicted to interact with at least three positions in the helix (**Extended Data Fig. 9k**). 136 of 227 (60%) profiled missense variants at the switch II helix increased KRAS abundance (**Fig. 6e,f**), an effect not observed in a previous mutational scan in yeast^82^. We hypothesized that the increased abundance of switch II helix variants could result from disruption of interactions with upstream GEFs such as SOS1, which destabilize the nucleotide-bound state, or with LZTR1, a CUL3 adapter protein that mediates KRAS ubiquitination and degradation, or from disruption of both, since both partners engage interfaces centered on the switch II helix (**Fig. 6e**)^83,84^.

To determine which interface positions form functional interactions with LZTR1 and SOS1, we repeated KRAS variant abundance assays in three additional cellular contexts: LZTR1-knockout cells, cells with elevated SOS1 activity, and both conditions combined (**Fig. 6f, Extended Data Fig. 10a,b**). Compared to basal conditions, the abundance of WT KRAS increased in LZTR1-knockout cells, indicating that LZTR1 promotes KRAS degradation (**Fig. 6g, Extended Data Fig. 10c**). Conversely, elevated SOS1 activity reduced the abundance of WT KRAS in both LZTR1-expressing and LZTR1-knockout cells, consistent with GEF interactions destabilizing KRAS (**Fig. 6g, Extended Data Fig. 10c**). By identifying where variant abundance deviated most from that of WT KRAS in LZTR1-knockout cells and in cells with elevated SOS1 activity, we resolved the positions that modulate each interaction (**Fig. 6f**, **Extended Data Fig. 10d**). 17 of the 22 KRAS positions at the LZTR1 interface were LZTR1-modulating, and 18 of the 32 at the SOS1 interface were SOS1-modulating, indicating that KRAS forms a similar number of functional interactions with each partner despite the difference in interface size (**Fig. 6h**, **Extended Data Fig. 10f**). Within each interface, interaction-modulating positions were split nearly evenly between partner-specific and shared ones. The eight shared positions were mainly in the switch II helix, and, at these positions, variant abundance deviated most from that of WT KRAS in response to both LZTR1 knockout and elevated SOS1 activity (**Fig. 6e,f, Extended Data Fig. 10e**). A further 11 SOS1-modulating positions, most of which bordered the SOS1 interface, and 14 LZTR1-modulating positions, most of which formed a network surrounding and extending from the β-phosphate of GDP, lay outside the interfaces. Structures could not have nominated these distal positions, which appear to allosterically modulate each interaction (**Extended Data Fig. 10f,g**).

To determine whether variants at non-interface positions affected abundance by altering LZTR1 engagement or by another mechanism, we used LABEL-seq to directly measure LZTR1 interaction with all 3,685 KRAS variants (**Extended Data Fig. 10h**). Variants at most non-interface LZTR1-modulating positions also altered KRAS–LZTR1 interaction (**Fig. 6i, Extended Data Fig. 10i**), indicating that these positions support presentation of the interface for binding. Our interaction data also shed light on the four LZTR1-modulating positions where variants paradoxically increased LZTR1-mediated degradation. At interface positions E62 and D105, variants increased interaction, promoting degradation by enhancing LZTR1 engagement (**Fig. 6i**). At non-interface positions E3 and E76 they did not increase binding, instead appearing to improve KRAS as a ligase substrate. Therefore, directly measuring KRAS-LZTR1 interaction validated the importance of interaction-modulating, non-interface positions and revealed mechanisms by which abundance-modulating variants act.

Intersecting structures with variant effects at scale assigned possible functional importance to 46 of 53 (87%) experimental and 324 of 361 (90%) predicted protein pairs. Abundance and pathway activity readouts yielded a variant-resolution map of disrupting each interface contact, separating interfaces where disruption increases target abundance while suppressing signaling, as at the RAF dimer interface, from those where disruption reduces abundance and signaling, as at RAF interfaces with HSP90 and CDC37. Iterating LABEL-seq across cellular contexts in which interaction partners are perturbed, alongside measuring KRAS-LZTR1 interaction, resolved the overlapping KRAS-LZTR1-SOS1 interface into partner-specific and shared binding determinants, revealed allosteric networks supporting each interaction, and uncovered distinct mechanisms by which variants promote LZTR1-mediated degradation. Thus, deep contextual interaction profiling resolved variant effect mechanisms that prior assays, structures, and variant effects measured in any single context could not.

## Discussion

We measured 503,099 variant effects across 17 proteins in the MAPK signaling pathway by pairing svClone, an efficient workflow for constructing comprehensive variant libraries of full-length human proteins, with LABEL-seq assays for protein abundance and pathway activity. Previous massively-scaled efforts focused on small protein domains and captured folding stability or surrogate functional outputs in cell-free or yeast models^8,9^. Smaller-scale efforts focused on individual MAPK pathway members and used transcriptional reporter or proliferation readouts, mostly in mammalian cells^36–52,85,86^. Our work bridges these two regimes, capturing multiple direct readouts of full-length MAPK protein function at unprecedented scale, thereby connecting intrinsic protein properties such as domain folding and allostery to complex phenotypic outputs in a human cellular context.

For example, pathway context transforms individual protein activities into pathway activity, which, when measured, revealed 14,412 dominant negative variants across 12 targets. These variants are depleted from the human population, demonstrating how at-scale knowledge of variant pathomechanism informs our understanding of selection in the human genome and hinting that dominant negative variants may be far more common than previously appreciated. Chaperone activity transforms protein stability into abundance inside the cell. Perturbing chaperone activity revealed that the abundance of ∼67% of missense variants depended on HSP90 activity, showing how variants that appear abundance-neutral under basal conditions can nevertheless create latent dependence on the cellular chaperone machinery^87^. Thus, while pervasive dominant negative variants impose an evolutionary constraint on signaling proteins, HSP90 relaxes a different constraint by sustaining variation that would otherwise reduce abundance. Neither constraint is apparent from sequence or structure, but pathway-scale variant effect data across cellular contexts revealed their contributions. The presence and activity of interacting proteins govern the effect of variants at interaction interfaces, including the overlapping KRAS switch II helix interface engaged by SOS1 and LZTR1, where variant effects depend on which partner is present. Thus, we demonstrate that the effect of most variants can only be understood as the product of their intrinsic effects on the target protein and the way these intrinsic effects are shaped by context.

The power of svClone and LABEL-seq to map variant effects at pathway scale comes with limitations. First, variants were expressed transgenically, so expression levels do not perfectly recapitulate those of the native protein. This limitation is particularly relevant for dominant negative variants, whose effects depend on the stoichiometry between the variant and endogenous pathway components. Second, all measurements were made in HEK293 cells under largely unstimulated conditions, capturing only one configuration of the pathway. Because signaling proteins adopt distinct regulatory states depending on the level and mode of pathway activation, variant effects that manifest only upon stimulation or in other cellular backgrounds were not detected. Third, we used phosphorylated ERK as a proxy for pathway activity. While ERK activity is the terminal readout of the pathway, it integrates upstream events and cannot distinguish among the many ways a variant might reshape signaling.

Our approach could be applied to other signaling pathways, and could be expanded to measure additional outputs such as alternative effector branches, transcriptional responses, and signaling dynamics. Pathway-scale variant effect mapping could shed light on additional druggable protein-protein interactions and could be combined with pharmacologic perturbations to reveal genotype-by-drug interactions and variant-specific therapeutic dependencies. Our work represents a step-change in the scale of variant effect mapping in human cells, illustrating the need to understand variant effects in their cellular context and considerably reducing the time and resources required to assemble a comprehensive atlas of variant effects^88^.

## Acknowledgements

We thank Andrew Stergachis, Stan Fields, Devin Schweppe and Christine Queitsch for helpful discussions and guidance. These results are in part based upon data generated by the TCGA Research Network: https://www.cancer.gov/tcga. We acknowledge the All Of Us Research Program participants, contributing institutions, and staff, without whom this research would not have been possible. This work was supported by the National Institutes of Health (RM1HG010461 and UM1HG011969 to D.M.F. and L.M.S., R35GM152106 to D.M.F., R01GM145011 and R01GM086858 to D.J.M., and R37CA252050 to A.H.B.) and the Chan Zuckerberg Initiative (CZIF2024-010284 to D.M.F. and L.M.S.). S.O. is a Washington Research Foundation postdoctoral fellow.

## Author Contributions

J.J.S., S.P., D.M.F. and D.J.M. conceived the study. J.J.S., S.P., A.F.A., M.W., Z.R.W., M.K.W.,

D.M.F. and D.J.M. developed the methodology. J.J.S. and S.P. developed software. J.J.S., S.P., A.F.A., M.W., A.J.V., Z.R.W., M.K.W., J.L.O., A.S. and M.H. performed experiments. J.J.S., S.P.,

A.F.A., M.W., Z.R.W., M.K.W. and A.F.R. performed validation. J.J.S., S.P., A.E.M. and S.O. performed formal analyses. J.J.S., S.P., A.F.A., M.W., Z.R.W. and A.S. provided resources. D.L.H., N.S. and A.F.R. curated data. J.J.S., S.P., D.L.H., D.M.F. and D.J.M. made the figures.

D.M.F. and D.J.M. managed the project. A.B., L.M.S., D.M.F. and D.J.M. supervised the work and acquired funding. J.J.S., S.P., A.F.A., Z.R.W., M.K.W., D.M.F. and D.J.M. wrote the manuscript. All authors reviewed and edited the manuscript.

---

**Extended Data Figure 1.**
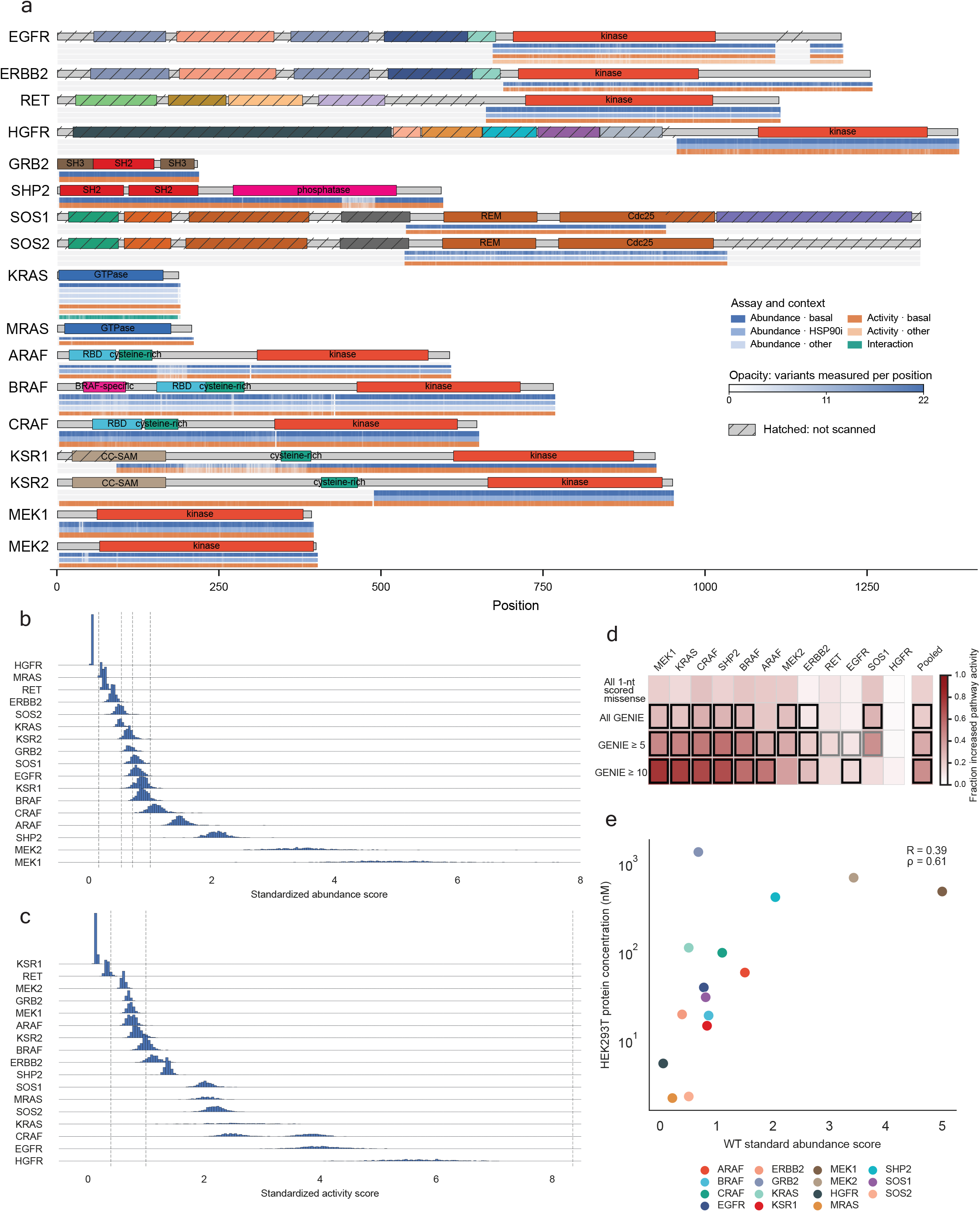
MAPK target protein domain structure and LABEL-seq assay dynamic range. **(a)** Schematic of domain architectures and variant coverage, drawn to scale by amino-acid position. Domain boundaries and names are from the NCBI Conserved Domain Database (accessed Aug 20, 2025); colored segments denote annotated domains, gray denotes unannotated sequence. Hatching denotes positions that were not varied in programmed libraries. Below each domain is a position-level plot showing the fraction of 22 missense, single-position deletion, or nonsense variants per protein measured in each assay and context. **(b, c)** Ridgeline plots of standardized **(b)** abundance and **(c)** pathway activity scores for synonymous variants of each of the 17 profiled target proteins under basal conditions, ranked by median standardized score. Dashed vertical lines mark reference scores. **(d)** Heatmap of the proportion of single-nucleotide missense variants classified as increased pathway activity under basal conditions for the targets meeting different tumor count thresholds, plus a pooled column combining all variants in listed targets (rightmost, separated by a gap; includes the five proteins — GRB2, KSR1, KSR2, MRAS, SOS2 — excluded from the individual protein columns for lacking such variants). Rows, top to bottom: proportion of increased pathway activity single-nucleotide missense variants in each target; proportion that are increased pathway activity among single-nucleotide missense variants observed at least once, observed at least five times, or at least ten times in the AACR Project GENIE tumor-sequencing registry. Box outlines denote cells with significantly more increased activity variants in the GENIE database as compared to all single-nucleotide missense variants (one-sided Fisher’s exact test p < 0.05, gray outline; p < 0.01, black outline). **(e)** Standardized abundance score versus steady-state HEK293T protein concentration (nM) for each profiled WT target^14^. Pearson R = 0.39; Spearman ρ = 0.61.

**Extended Data Figure 2.**
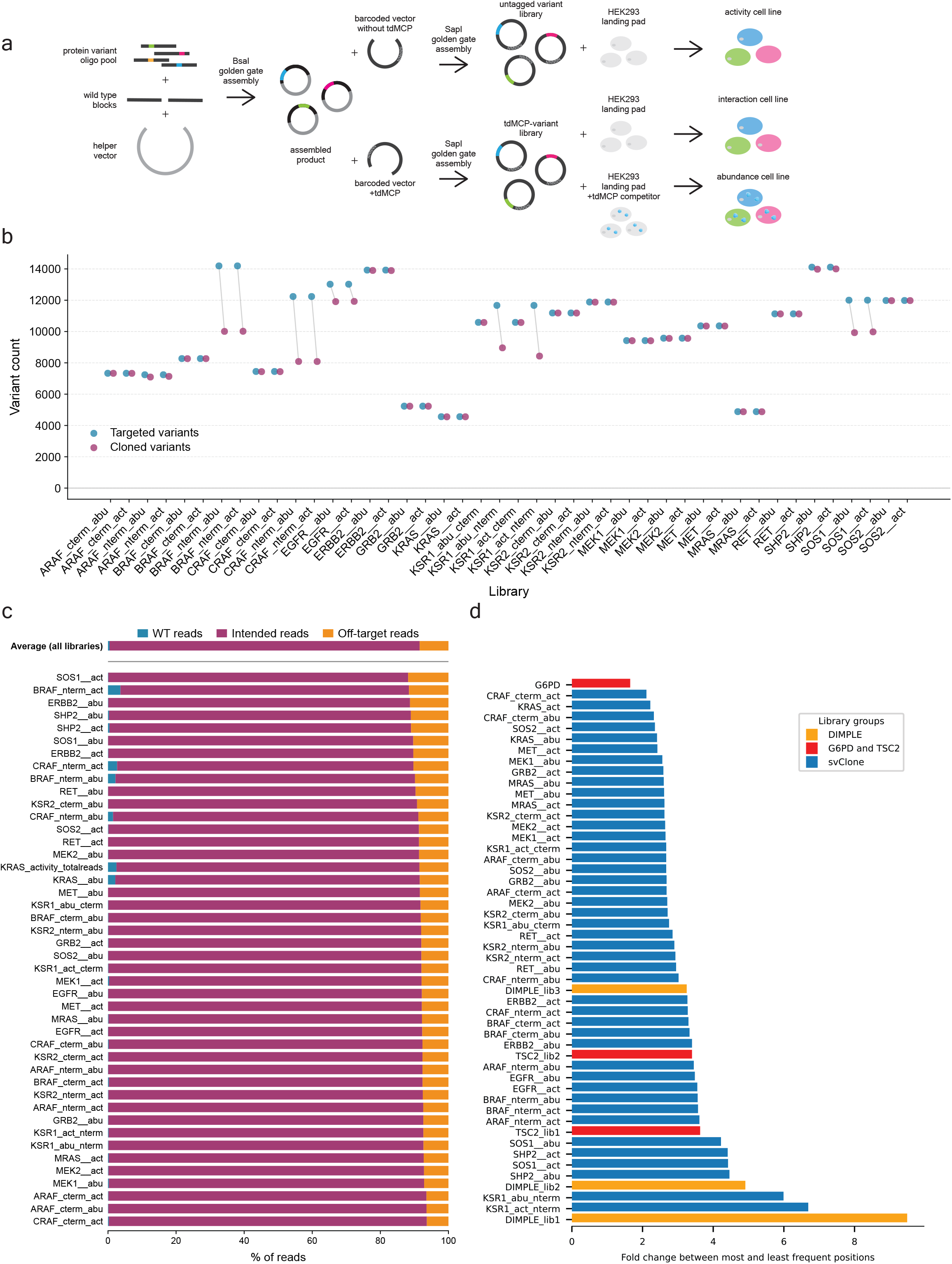
Synthetic DNA variant cloning (svClone) workflow and performance. **(a)** svClone workflow, with tdMCP tagging scheme. svClone employs array-synthesized oligonucleotide pools encoding all single amino acid substitutions, 3-nucleotide deletions and selected synonymous and nonsense variants, assembled with invariant synthetic DNA blocks into full-length open reading frames. **(b)** Number of programmed variants vs number of variants successfully cloned and paired with at least one barcode for each of the 44 activity (act, untagged variants) and abundance (abu, Flag-tdMCP-tagged) libraries used in this study. Programmed variants are the total number of unique designed variants in the synthesized oligo pool (synonymous, missense, nonsense, deletions). Gray lines connect the paired values for each library. Recovery (cloned/targeted variants) ranged from 66.1% to 100% across libraries (mean 94.6%). HGFR libraries are labeled as MET. **(c)** For each library, the fraction of sequencing reads from the final libraries corresponding to wild type (WT), programmed variants, or non-programmed variants. Across libraries, programmed variants accounted for 84.8–93.5% of reads (mean 91.1%), non-programmed variants for 6.3–11.8% (mean 8.4%), and WT for 0.03–3.65% (mean 0.45%). Among all 202,520 observed programmed variants, 195,204 had ≥5 barcodes per variant in either pathway activity (untagged) or abundance (tdMCP-tagged) libraries. **(d)** Barplot showing position-level skewness across libraries, expressed as the fold-change between the most and least frequently occurring positions. Libraries from a commercial source (red)^92,93^ and DIMPLE, another array-synthesized oligonucleotide pool-based methods (gold)^12^, are shown for comparison.

**Extended Data Figure 3.**
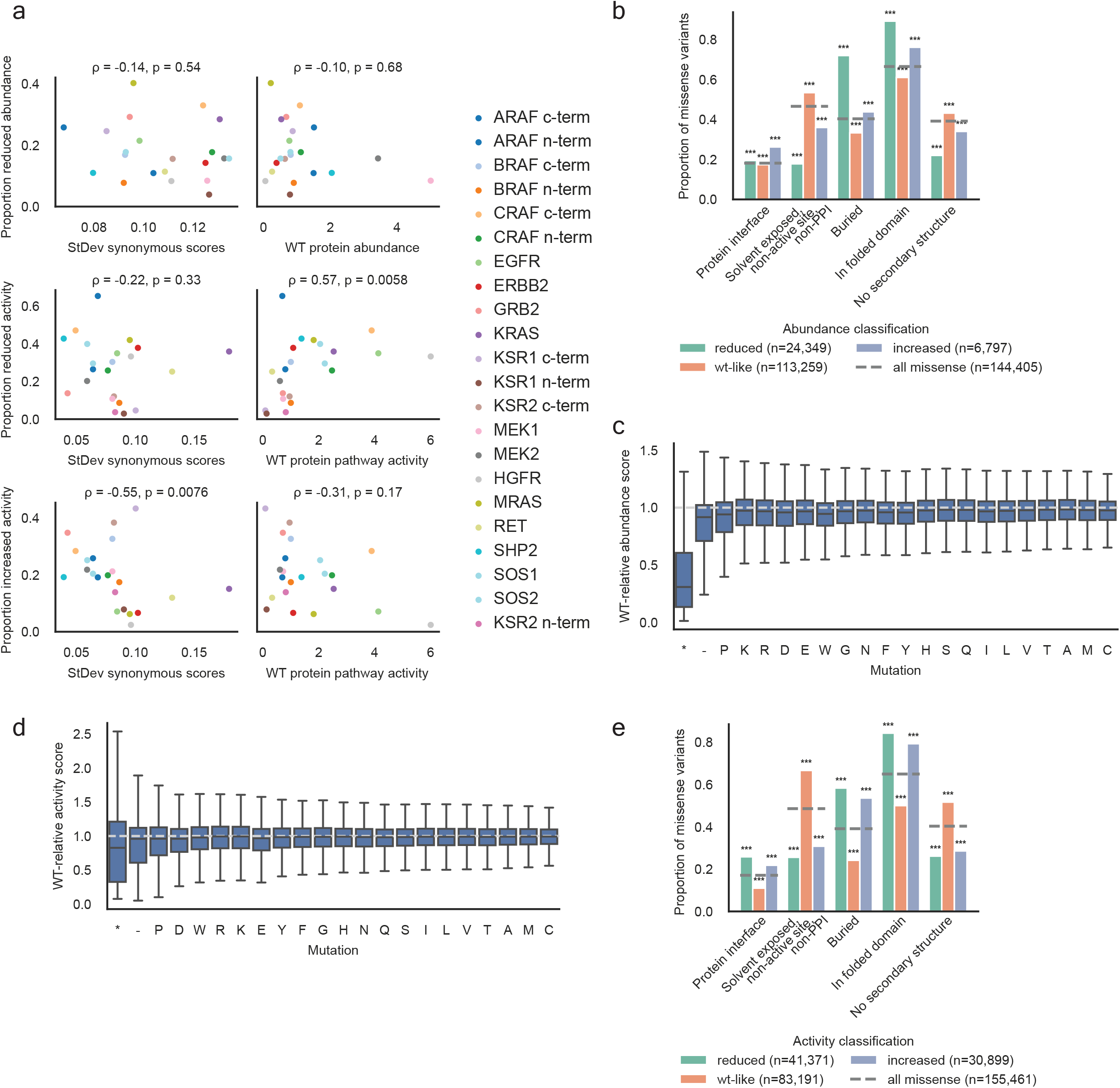
Summary of pathway-scale MAPK variant abundance and pathway activity data. **(a)** Relationship between per-library quality metrics and the proportion of missense variants classified as reduced or increased abundance or activity under basal conditions. Each point is one library (target, or N-/C-terminal sub-library for multi-library proteins). Top row: proportion of missense variants classified as reduced abundance versus the standard deviation of synonymous WT-relative abundance scores (left) and versus the wild type (WT) standardized abundance score (right). Middle row: as in top row, for the proportion classified as reduced pathway activity. Bottom row: as in top row, for the proportion classified increased pathway activity. Spearman’s ρ and p-value from a two-sided Spearman rank correlation test shown. **(b)** Proportion of missense variants in each abundance class, grouped by structural feature. Dashed gray line, proportion among all scored missense variants. Asterisks denote a permutation test p-value for variants in the indicated abundance class versus all variants (100 group-size-matched resamples). *** indicates p-value<0.001. **(c)** Distribution of WT-relative abundance scores for each amino acid substitution, pooled across all 17 targets, with single-position deletion and nonsense variants shown for comparison. Box, 25th–75th percentile; whiskers extend to the most extreme non-outlier value. n = 155,917 score measurements. Dashed line, WT score. **(d)** As in (c), for WT-relative pathway activity (n = 171,149 score measurements). **(e)** As in (b), for different pathway activity classifications.

**Extended Data Figure 4.**
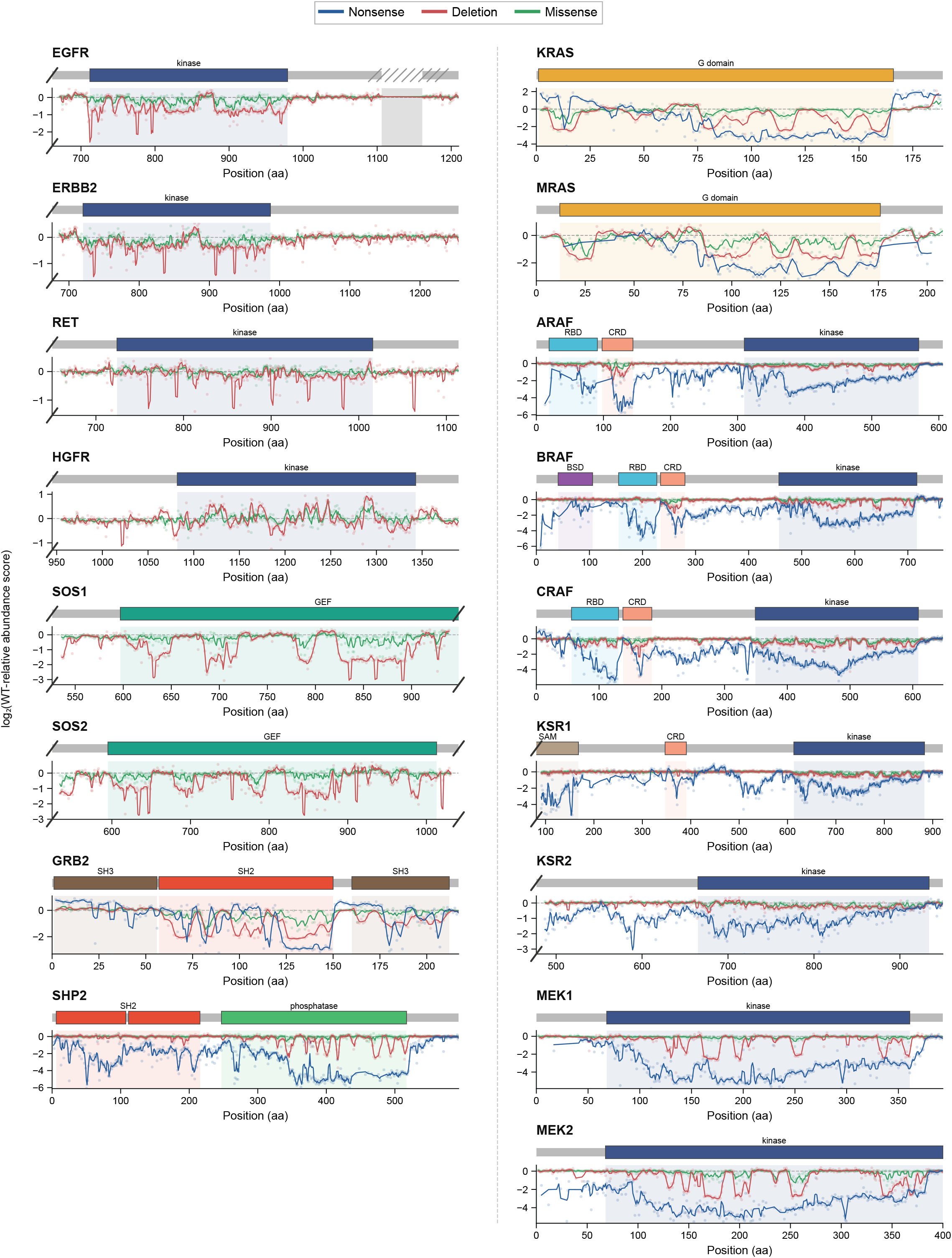
Abundance scores and domain architecture across all target proteins. For each target, top panel shows domain annotations for scored positions, with diagonal hash marks indicating unscored regions. Bottom panel shows position mean WT-relative abundance scores, colored by variant type under basal conditions. Line denotes a smoothed 3-position rolling mean; individual position means are shown as points. Dashed line denotes the wild type abundance score.

**Extended Data Figure 5.**
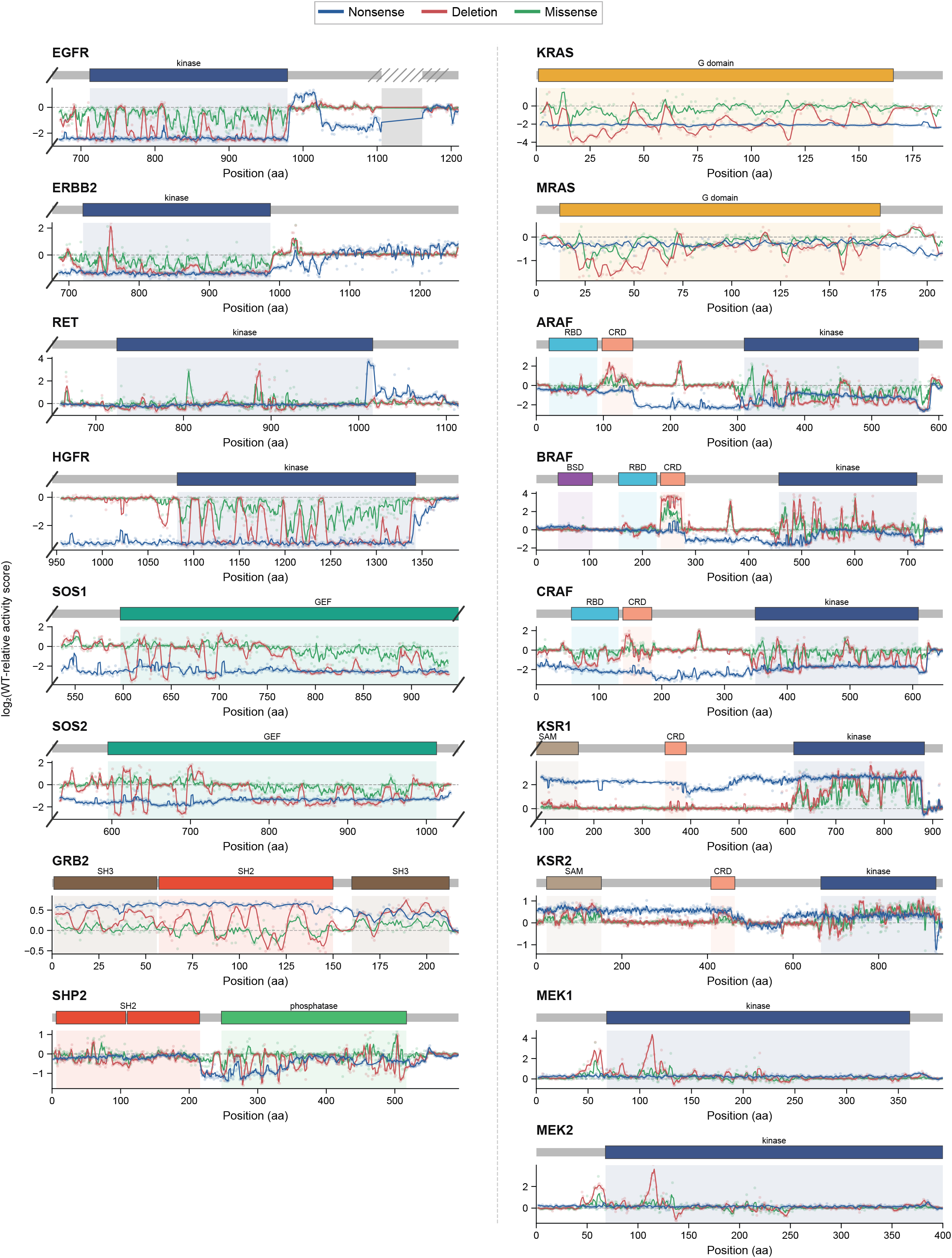
Pathway activity scores and domain architecture across all target proteins. For each target, top panel shows domain annotations for scored positions, with diagonal hash marks indicating unscored regions. Bottom panel shows position mean WT-relative pathway activity scores, colored by variant type under basal conditions. Line denotes a smoothed 3-position rolling mean; individual position means are shown as points. Dashed line denotes the wild type pathway activity score.

**Extended Data Figure 6.**
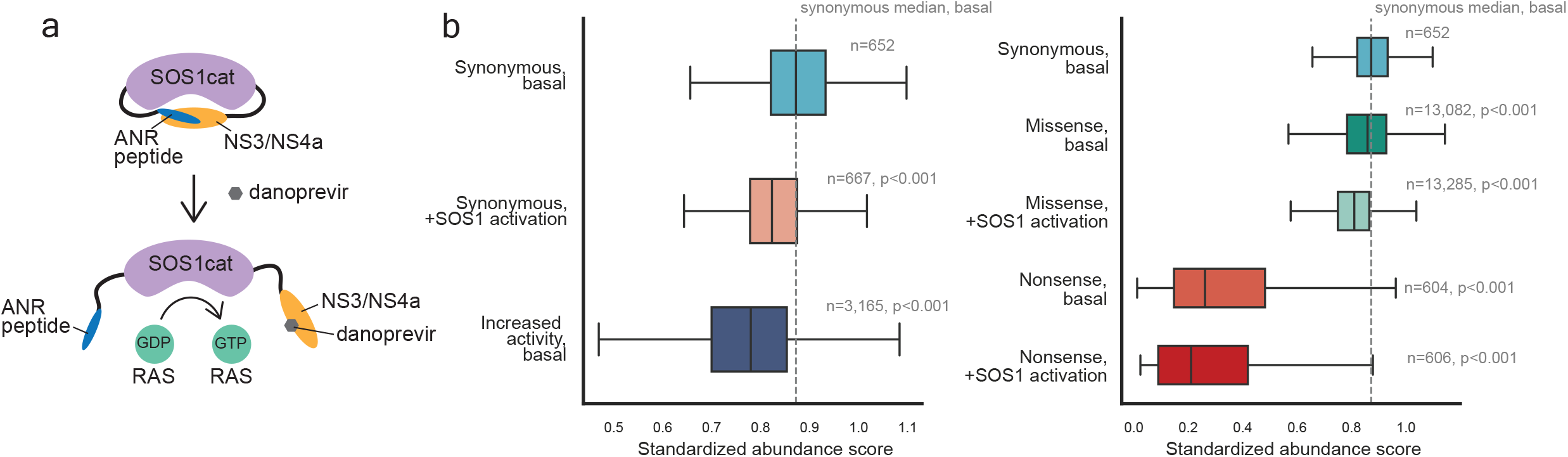
Relief of autoinhibition reduces BRAF abundance. **(a)** Schematic of the chemically inducible activator of RAS (CIAR) switch, comprising a constitutively active RAS-activating module of the GEF SOS1 that is autoinhibited by the interaction between NS3a and the peptide ANR. Treatment of cells with the bioorthogonal NS3a-binding drug danoprevir relieves SOS1 autoinhibition, raising intracellular RAS-GTP levels. Elevated RAS-GTP levels disrupt the autoinhibited state of BRAF by promoting its recruitment to active signalling complexes at membranes. See Methods for use of CIAR in the variant-library abundance and activity assays^94^. **(b)** Standardized BRAF abundance scores for the indicated variant types under basal conditions or under the elevated KRAS-GTP levels produced by CIAR. Box, interquartile range; center line, median; whiskers extend to the most extreme non-outlier value. Dashed vertical line, median basal synonymous wild type score. P-values, two-sided Mann-Whitney U test comparing each group against the basal synonymous wild type distribution.

**Extended Data Figure 7.**
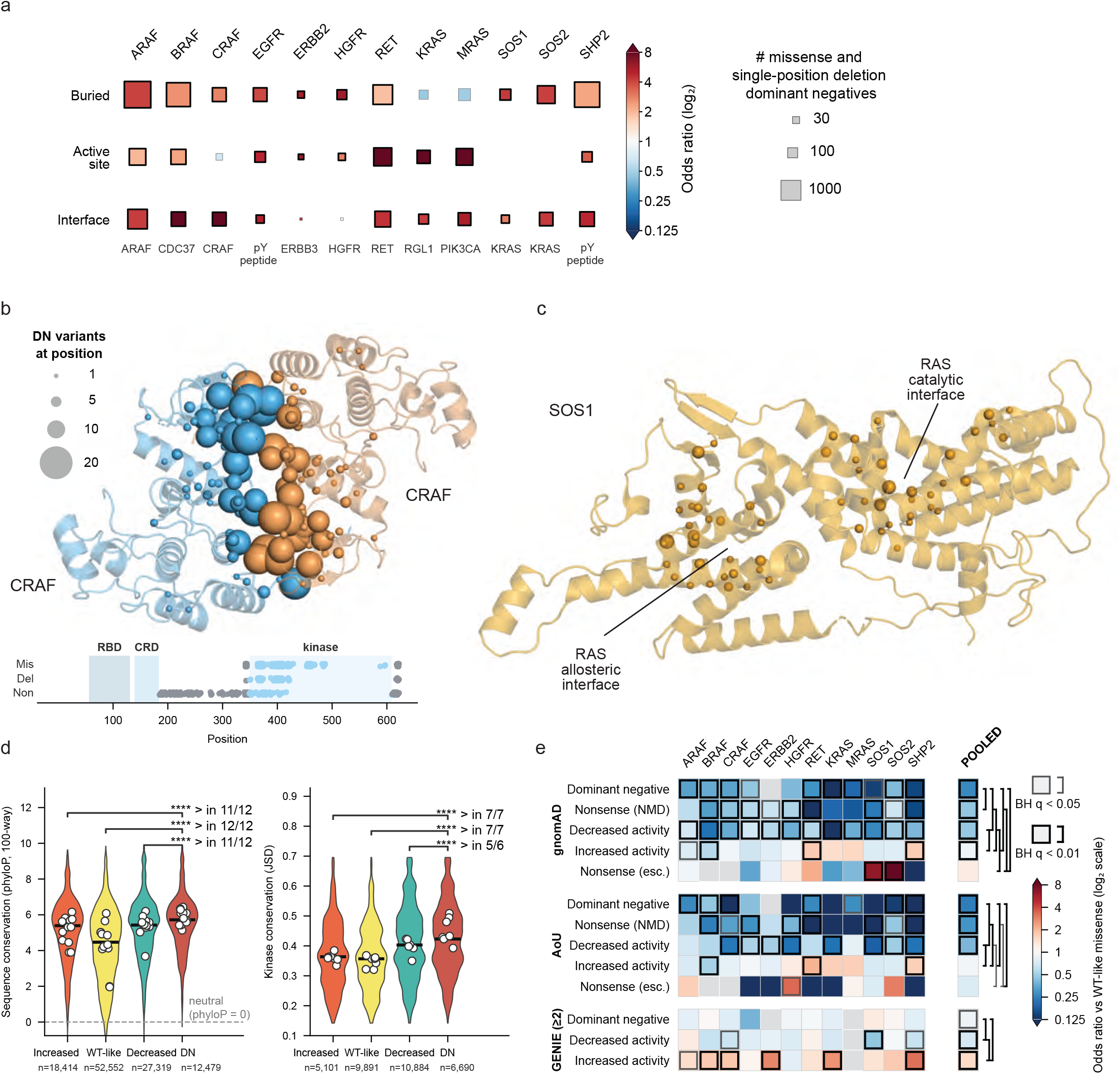
MAPK pathway dominant negative variant biochemical features, conservation and depletion in the human population. **(a)** Enrichment or depletion of pooled missense and single-position deletion dominant negatives at buried, non-active site, active site, or interface positions in each target (see Methods). Size of box indicates number of dominant negative variants and color indicates odds ratio for dominant negative in each category versus non-dominant negative. Under the interface boxes the name of the most enriched interaction partner is shown. Black border indicates significance (Fisher’s exact test, q < 0.01). **(b)** Dominant negatives shown on CRAF kinase domain dimer (PDB 8CPD). Mutations at the dimer interface are enriched for dominant negatives (Fisher’s exact test, q < 0.001). Missense, single-position deletion and nonsense (truncating) dominant negatives shown below by location in protein. **(c)** Dominant negatives mapped onto the structure of SOS1 bound to HRAS (PDB 1NVU), with sphere sizes as in (b). The catalytic and allosteric RAS-binding interfaces are labelled; HRAS not shown, for clarity. **(d)** Distributions of 100-way phyloP scores for all protein positions (left) or kinase conservation scores (see Methods) for kinase domain positions (right), for positions containing variants in each pathway activity class. White dots represent per-protein medians. *** indicates significance (Mann-Whitney U-test, q < 0.001) and > indicates number of proteins where median value is higher in that comparison. **(e)** Enrichment or depletion of missense variants by pathway activity class, relative to WT-like activity variants, for each protein or pooled (column on right) in gnomAD (top), the AllOfUs biobank (AoU, middle), or GENIE catalog of somatic variation in cancer (bottom). Nonsense (NMD) indicates nonsense variants >50 nucleotides upstream of the last exon-exon junction codon, and therefore subject to nonsense-mediated decay when expressed endogenously. Nonsense (esc.) indicates nonsense variants within 50 nucleotides of the last exon-exon junction, which escape nonsense-mediated decay when expressed endogenously. Color indicates odds ratio for variants in the indicated activity class versus variants with WT-like activity; gray or black border indicates significance of enrichment or depletion versus variants with WT-like activity; gray or black brackets indicate pairwise significance of enrichment or depletion between activity classes (Fisher’s exact test, q < 0.05 or < 0.01 respectively).

**Extended Data Figure 8.**
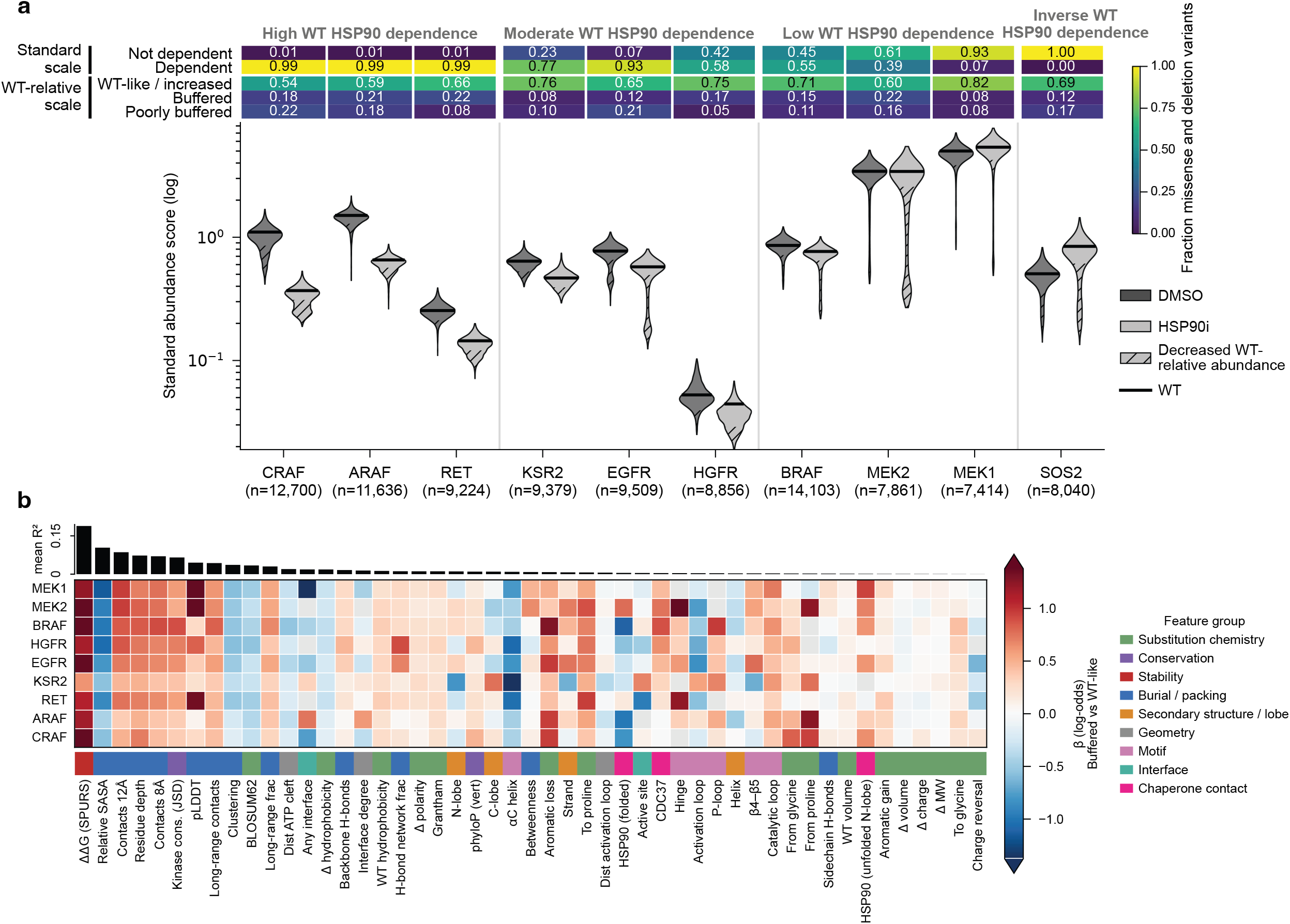
Variant effects under HSP90 inhibition and feature analysis. **(a)** Summary of basal and HSP90 inhibited (HSP90i) missense and 3-nucleotide deletion abundance scores across all positions within each target. Top: heatmap showing the fraction of variants that are dependent or not dependent on HSP90 (standardized scale) and buffered, poorly buffered, or WT-like/increased (WT-relative scale). Colors indicate the fraction of variants in each category. Bottom: violin plots showing abundance score distributions for each target, grouped by WT HSP90 dependence. Bars indicate the WT score, violins are trimmed to the 0.25th–99.75th percentile for display. Hatching indicates the portion of the variant distribution that is low abundance (see Methods for classification). **(b)** Heatmap of univariate logistic-regression coefficients (β, log-odds) for buffered versus WT-like kinase-domain missense variants, fit separately for each (protein, feature) pair on z-scored feature values. Red indicates higher feature values are associated with buffering; blue indicates association with WT-like/high (unbuffered) abundance.

**Extended Data Figure 9.**
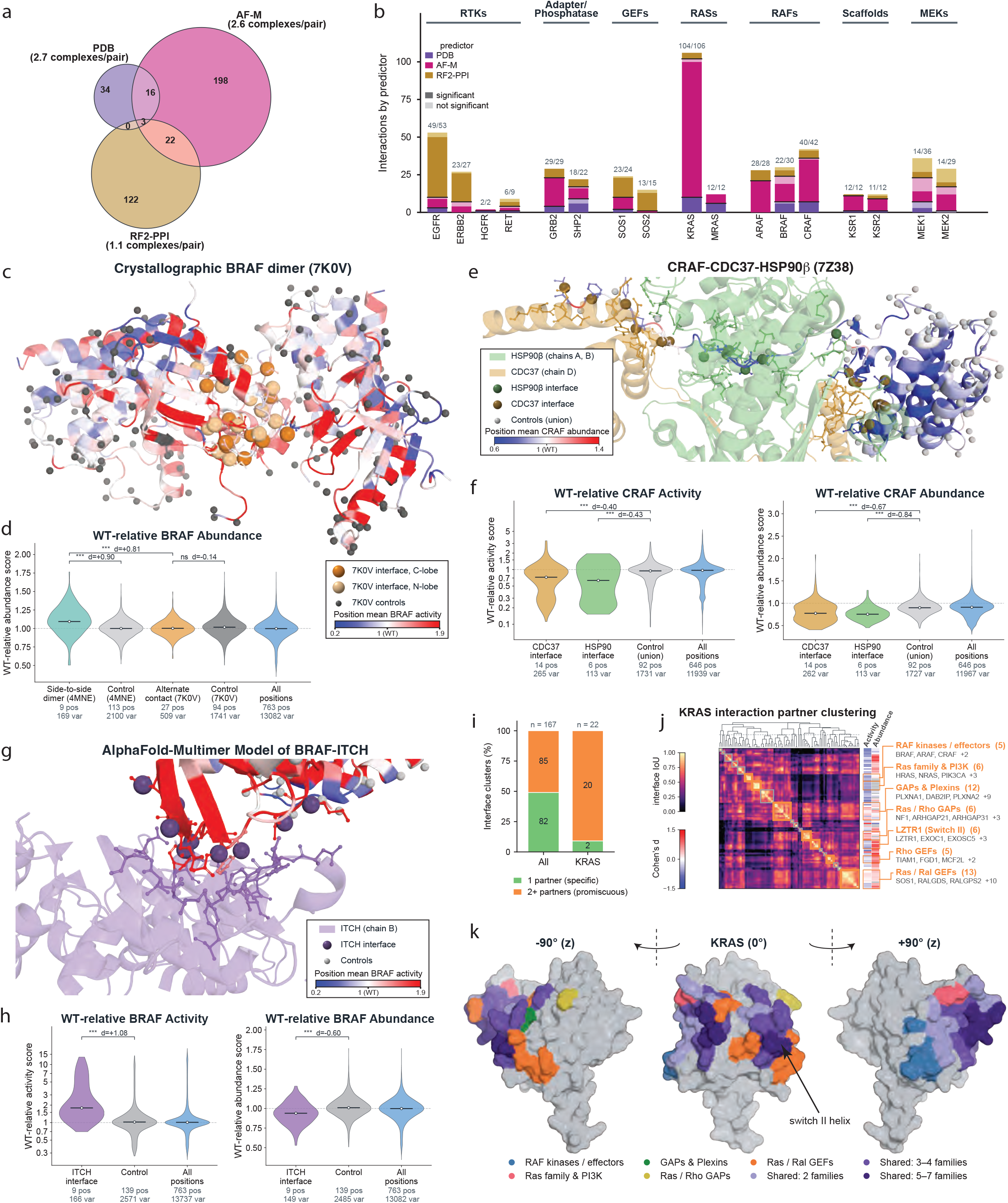
Systematic analysis of variant effects at MAPK pathway target interaction interfaces. **(a)** Venn diagram showing the number of unique protein pairs with at least one assessable complex from the PDB or predicted by AF-M or RF2-PPI. **(b)** Unsupported (light shading) and functionally supported (dark shading) interaction partners of each scored protein, colored by origin (PDB versus predictor) and organized by protein function. In this plot, a protein pair is supported if at least one of its complexes from the indicated source shows a significant difference in either pathway activity or abundance scores at the interface compared to control positions (see Methods, Fig. 6a), and is counted once per source that models it. Above each bar is the supported / total number of protein pairs. **(c)** BRAF WT-relative activity scores mapped onto an artifactual crystallographic dimer structure (PDB 7K0V). Interface positions in each protomer are shown with ball-and-stick and orange spheres, shaded by protomer; control positions are indicated with gray spheres. **(d)** BRAF WT-relative abundance score distributions for positions at the canonical activating dimer (PDB 4MNE) and the artifactual crystallographic dimer (PDB 7K0V) interfaces, each compared with its own control positions, and for all positions. Brackets indicate statistical comparisons (*** indicates Mann-Whitney U-test FDR-corrected q < 0.001), Cohen’s d value shown for comparison, and the number of variants (var) and positions (pos) are indicated below each group of positions. **(e)** CRAF WT-relative basal abundance scores mapped onto the CRAF-CDC37-HSP90β structure (PDB 7Z38). Interface positions on CRAF and on the interactors are shown as ball-and-stick, and spheres mark CRAF interface and control positions with color denoting interactor or control. **(f)** CRAF WT-relative activity (left) and basal abundance (right) score distributions for positions that are part of CDC37 or HSP90β interfaces (PDB 7Z38), each compared with a shared control set of positions, and for all positions. Brackets as in (d). **(g)** BRAF WT-relative activity scores mapped onto an AF-M model of the BRAF-ITCH complex, with ITCH shown in purple; ITCH positions that interact with BRAF are shown as sticks. BRAF interface positions are shown as ball-and-stick, and light grey spheres indicate BRAF control positions. **(h)** BRAF WT-relative activity (left) and abundance (right) score distributions for BRAF positions that are part of the ITCH interface, control positions, and for all positions. **(i)** For each profiled target, interface positions of partners containing at least 5 scored positions were clustered using intersection-over-union distance (IoU, see Methods). Stacked bars show the number of resulting interface clusters containing a single partner (green) or by 2 or more partners (”promiscuous”, orange) for all proteins or for KRAS alone. **(j)** Heatmap for pairwise IoU of the interface positions of interaction partners of KRAS. The heatmap annotation on the right shows each partner’s median missense Cohen’s d over its complexes for activity and abundance. Gray indicates that all complexes at that interface had q > 0.05. Interface cluster names and key interaction partners at each interface cluster are shown, along with the number of interaction partners in the cluster in parentheses. **(k)** Three views of the KRAS surface related by 90° rotations about the vertical axis with interface positions colored by the clusters annotated in (j). A position is assigned to a cluster if at least 50% of that cluster’s partners contact it. Positions belonging to a single cluster are colored by that cluster and positions shared by at least 2 clusters are shaded by the number of clusters that contact them. The KRAS switch II helix is marked by an arrow.

**Extended Data Figure 10.**
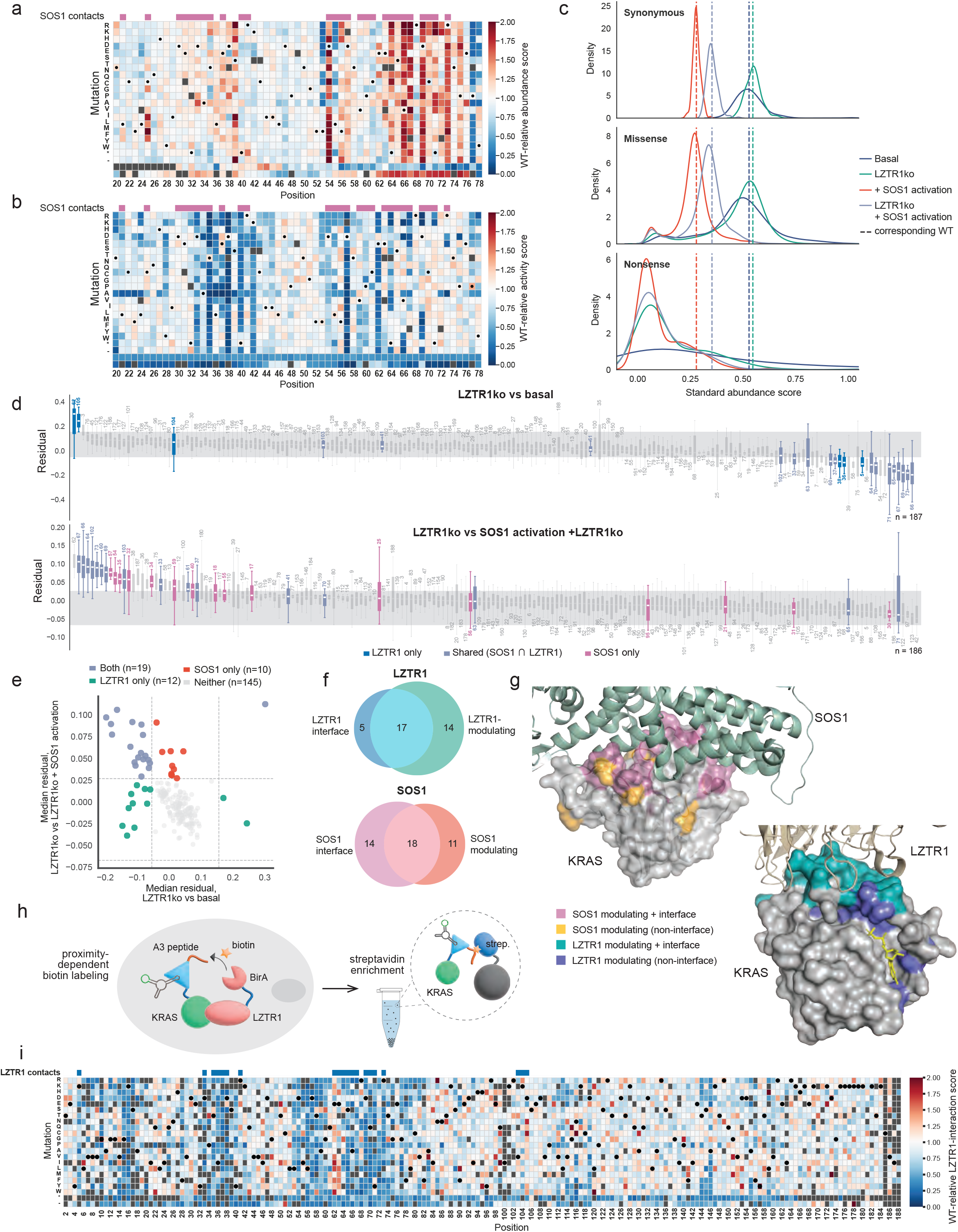
Dissection of KRAS-SOS1 and KRAS-LZTR1 interactions using LABEL-seq. **(a)** Variant effect map showing WT-relative abundance scores for all variants at KRAS positions 20–78 under SOS1 activation. Each column represents a single position and each row a substitution to the indicated amino acid. Dots indicate the WT amino acid; color indicates score; dark gray cells indicate missing data. SOS1 interface positions from Fig. 6e are shown above the map. **(b)** Variant effect map as in (a) showing the WT-relative pathway activity scores for all variants at KRAS positions 20–78 under SOS1 activation. **(c)** Density plots of standardized abundance scores for synonymous (top), missense (middle), and nonsense (bottom) KRAS variants under four conditions: basal (navy), LZTR1ko (teal), SOS1 activation (red), and SOS1 activation in LZTR1ko cells (lavender). Dashed vertical lines represent the WT score under each condition. **(d)** Box plots representing the distribution of per-variant residuals across all missense substitutions at a given KRAS position (box, Q1–Q3; horizontal line, median; whiskers, 1.5 × IQR). Residuals were derived from an origin-constrained linear regression of abundance scores between two conditions. Positions are ordered left to right by descending median residual. The gray shaded band indicates the 5th-95th percentile range of synonymous variant residuals; positions with median residuals outside this band are classified as modulating the LZTR1 or SOS1 interaction. Top: residuals from LZTR1ko versus basal conditions. KRAS positions at the LZTR1 interface (PDB 9MF0) are indicated: positions unique to the LZTR1 interface (blue) and positions shared with the SOS1 interface (lavender); positions unique to the SOS1 interface and non-interface positions are shown in gray. Bottom: residuals from SOS1 activation in LZTR1ko cells versus LZTR1ko. KRAS positions at the SOS1 interface (PDB 1NVU) are indicated: positions unique to the SOS1 interface (rose) and positions shared with the LZTR1 interface (lavender); positions unique to the LZTR1 interface and non-interface positions are shown in gray. **(e)** Scatter plot showing the relationship between LZTR1-modulating positions and SOS1-modulating positions. Each point represents one KRAS position, plotted by its median missense-variant residuals from (d). Positions are colored by their functional status: modulating both the LZTR1 and SOS1 interactions (lavender), LZTR1 interaction only (teal), SOS1 interaction only (red), or neither (gray). Dashed lines denote thresholds for modulating positions. **(f)** Venn diagrams of the overlap of LZTR1 interface positions (PDB 9MF0) with LZTR1 interaction-modulating positions (top) and of SOS1 interface positions (PDB 1NVU) with SOS1 interaction-modulating positions (bottom), as defined in (d). **(g)** KRAS surfaces (left, PDB 1NVU; right, PDB 9MF0) colored according to interface and non-interface LZTR1-interaction (left) or SOS1-interaction (right) modulation classification. **(h)** Schematic of the KRAS–LZTR1 interaction assay. BirA-LZTR1 is co-expressed with the tdMCP-KRAS variant library fused to the peptide A3, a weak substrate for BirA biotin ligase, which labels A3-tdMCP-KRAS variants in a proximity-dependent manner. In-cell biotin labelling followed by streptavidin enrichment yields variant scores reflecting KRAS–LZTR1 interactions. **(i)** Variant effect map showing the WT-relative interaction scores for all KRAS variants with LZTR1. Annotations as in (a).

## Supplementary Table Legends

**Supplementary Table 1 | Protein target descriptions.** Biochemical descriptions of all target proteins.

**Supplementary Table 2 | Variant scores and annotations.** LABEL-seq abundance, pathway activity, and interaction scores as well as annotations for all profiled variants.

**Supplementary Table 3 | Variant effects at interfaces**. Annotations and interface versus control statistics for every scored interface in a predicted or experimentally-determined structure.

**Supplementary Table 4 | Oligos.** Oligos used in cloning and preparation of amplicons.

**Supplementary Table 5 | Plasmids.** Description of plasmids and addgene accession numbers.

**Supplementary Table 6 | Reagents.** Reagent catalog numbers.

## Data, Code and Reagent Availability

### Data availability

All variant effect scores, raw sequencing reads and barcode-to-variant maps will be available on the IGVF portal (data.igvf.org, project accession: IGVFDS0431GNGK). Variant effect scores and annotations can also be found in Supplementary Table 2, and are available on MaveDB under accessions urn:mavedb:00001302-urn:mavedb:00001318^95^. Tabular data is provided as a supplement or, for larger files, is available via Zenodo at https://doi.org/10.5281/zenodo.23121366.

### Code availability

Code for designing variant libraries using svClone is available at github.com/FowlerLab/svClone. Code necessary to reproduce our analyses, starting at the raw sequencing reads or from summary data available via supplementary tables and Zenodo, can be found at github.com/MalyLab/LABEL-seq. Code for assembling variant-barcode maps from long read sequencing data is available at github.com/FowlerLab/pacybara.

### Material availability

Plasmids will be available on addgene (**Supplementary Table 5**). Other materials available upon request.

## Methods

### Miscellaneous

A list of oligo sequences and available plasmids are provided in **Supplementary Table 4** and **Supplementary Table 5**, respectively. A list of reagents and supplier catalog numbers is provided in **Supplementary Table 6**.

### Protein target selection and variant library design

We selected 17 proteins representing the core of the canonical RAS–MAPK signaling pathway on the basis of their clinical relevance, biochemical characterization, and suitability for overexpression in HEK293 cells. MANE-Select CDS sequences were pulled from GenBank to serve as the templates for mutagenesis, except for KSR1 which used UniProt Q8IVT5 (canonical isoform) and KRAS which used UniProt P01116-2 (KRAS4B). In some cases, synonymous mutations were introduced to eliminate recognition sites for restriction enzymes (BsaI, SapI, and NheI-HF) used in downstream cloning or codon optimization to allow for synthesis of gene blocks by IDT using the IDT Codon Optimization Tool (https://www.idtdna.com/CodonOpt). Each protein was computationally divided into varying numbers of tiles containing approximately 63 amino acids and appended with overhangs containing restriction sites for Golden Gate assembly cloning and retrieval primers (https://github.com/FowlerLab/svClone). We designed these tiled oligonucleotide pools to encode 19 single amino acid substitutions as well as synonymous variants, stop codons, and 3-nucleotide deletions in every frame at every position using a codon optimization strategy that minimized the number of nucleotide changes per variant. All oligos for each tile were scanned for new restriction sites that conflict with the cloning workflow and for the presence of duplicated sequences; identified oligos were eliminated from the pool prior to commercial synthesis (Twist Bioscience). Complementary cloned gene blocks with wild-type (WT) sequences were synthesized and sequence confirmed (Twist Bioscience, IDT) to allow for BsaI-HF (New England Biolabs) Golden Gate assembly of full length ORF libraries into intermediate vectors prior to cloning into barcoded assay vectors by SapI (New England Biolabs) Golden Gate Assembly (**Extended Data Fig. 2a**). To keep library cost as low as possible, given that cloned gene blocks increase in price past a certain length, proteins encoded by sequences longer than 1,800 bases required additional corresponding WT gene blocks to complete the full gene when assembled into each final assay vector using Golden Gate assembly by SapI.

### Cloning barcoded variant libraries

A total of four attB-MS2-circRNA assay vectors were used to generate variant library plasmids. For protein abundance and interaction profiling, two attB-MS2-circRNA plasmids containing 3×Flag-tdMCP tags were designed to fuse to the protein of interest at either the N-terminus or the C-terminus (**Supplementary Table 5**). For pathway activity profiling, a similarly designed attB plasmid was used which included the MS2-circRNA, but in which the target protein was not tagged with tdMCP (untagged).

#### Barcoding tdMCP and untagged assay vectors

Prior to SapI Golden Gate Assembly of variant libraries, a 16-nucleotide barcode was inserted into all attB plasmids. First, a ssDNA oligo (SP020924, 10 μM) containing a degenerate 16-nucleotide region was annealed with primer SP187 (10 μM, at 98°C for 3 min, followed by ramping down to 25°C at −0.1°C per second). After annealing, 1.35 μL (1 mM) dNTPs and 0.8 μL Klenow exo-polymerase (New England Biolabs) were added to fill in the barcode oligo. The reaction was placed in a thermocycler at 25°C for 15 min, 70°C for 20 min, then ramped down to 37°C in −0.1°C per second increments. Barcode fragments were visualized on an Agilent TapeStation 4200 (Agilent Technologies) to confirm that the size was correct. Duplicate reactions were pooled and cleaned using Zymo DNA Clean & Concentrator-5 (Zymo Research) following the protocol for PCR product cleanup, and eluted in 20 μL elution buffer. Barcode fragments were quantified using Qubit dsDNA HS Assay Kit, then used as the insert in a Golden Gate Assembly with Esp3I (New England Biolabs) at a 5:1 molar ratio of insert:vector. Each reaction consisted of: 2.5 μL T4 DNA Ligase Buffer, 0.5 μL T4 DNA Ligase, 1.0 μL Esp3I, 250 ng target vector, barcode fragment DNA, and nuclease-free water to bring the reaction volume to 25.0 μL total. Reactions were run using the following program: 30 cycles of 37°C for 5 min followed by 16°C for 5 min, then a heat-inactivation at 60°C for 15 min, another heat-inactivation at 85°C for 15 min, and a hold at 4°C. Upon completion of the program, products underwent an additional digest with BsmBI-v2 (New England Biolabs). The following was added directly to each reaction: 1 μL Buffer r3.1 (New England Biolabs), 1 μL BsmBI-v2, and 3 μL nuclease-free water. The additional digest was run on a thermocycler at 55°C for 2 h, followed by heat-inactivation at 80°C for 15 min, and a hold at 4°C. Reactions were cleaned using Zymo DNA Clean & Concentrator-5 following the protocol for plasmid cleanup. Cleaned, barcoded products were quantified using Qubit dsDNA HS Assay Kit. These barcoded products were transformed into NEB 10-beta Electrocompetent *E. coli* (New England Biolabs) in replicate 25 μL reactions according to the manufacturer’s protocol using a BioRad Gene-Pulser electroporator set at 2.0 kV. The recovery cultures (2 mL) were plated on LB-ampicillin 24.5 cm x 24.5 cm square plates, and grown overnight at 30°C. Dilution plates were used to calculate the number of transformants and estimate the number of barcodes. Square plates were harvested using cold LB broth and cells were gently scraped from the plate surface and transferred to a 50 mL conical tube. Once the entire plate was harvested, the bacteria were centrifuged at 4,000 x *g* for 20 min at 4°C. Barcoded plasmid DNA was isolated using the ZymoPURE II Plasmid Maxiprep Kit (Zymo Research) according to manufacturer’s instructions. Plasmid preps were quantified using Qubit dsDNA BR Assay Kit.

#### Oligo retrieval

Each subpool of variant-encoding oligos corresponding to a tile was retrieved from the synthesized oligo pool via real-time qPCR using pre-programmed orthogonal forward and reverse primers^96^. The oligo pool was diluted to 1 ng/μL and used as the template for a reaction containing 25 μL Q5 Hot Start High Fidelity 2X Master Mix (New England Biolabs), 0.3 μL SYBR green I (ThermoFisher Scientific), orthogonal retrieval primers^96^ (RR_orthogonalF/Rv1_1-82, **Supplementary Table 4**) at a final concentration of 0.5 μM, and nuclease-free water to bring the total volume to 50 μL per reaction. Samples were run on a CFX Connect (Bio-Rad) for approximately 21 cycles or until all samples reached 3,000 relative fluorescence units (RFU). Reactions were denatured at 98°C for 30 seconds, followed by cycles of 98°C for 30 seconds, 63°C for 10 seconds, and 72°C for 20 seconds. Products were cleaned using 1.5X SeraMag beads (Cytiva) and eluted into 32 μL of nuclease-free water, following the manufacturer’s protocol. Cleaned products were quantified using a Synergy H1 Hybrid plate reader (BioTek). To quantify variant representation, a dilute portion of retrieved oligos were then amplified for 10 cycles using Nextera XT indexing primers and sequenced on an Illumina NextSeq 2000 instrument using a NextSeq 1000/2000 P2 300 Cycle kit (Illumina).

#### Assembly of amplified variant oligo tiles into helper vector

Next, each amplified tile was assembled with gene blocks completing the ORF into pHSG299-BsaI, an intermediate vector modified from pHSG299 (Takara 3299) to contain BsaI recognition sites for Golden Gate assembly. For each tile, duplicate Golden Gate reactions comprising 2.5 μL T4 DNA Ligase Reaction Buffer (New England Biolabs), 0.5 μL T4 DNA ligase (New England Biolabs), 1.0 μL BsaI-HFv2, 100 ng pHSG299-BsaI plus an appropriate amount of insert DNA (i.e. amplified tile and 1-2 gene blocks at a 2:1 insert:vector molar ratio), and nuclease-free water to bring the total volume to 25 μL. Reactions were run in a thermocycler at 37°C for 18 h, followed by a heat inactivation at 80°C for 10 min and stored at 4°C. After the reactions were completed, products underwent an additional digest with BsaI-HFv2 and Exonuclease V (New England Biolabs). The following was added directly to each reaction: 0.25 μL BsaI-HFv2, 0.25 μL Exonuclease V, 0.5 μL 10X rCutSmart Buffer (New England Biolabs), and 4 μL nuclease-free water. The additional digest was run on a thermocycler at 37°C for 2 h, followed by heat-inactivation at 80°C for 10 min, and a hold at 4°C. Duplicate reactions were pooled and cleaned using Zymo DNA Clean & Concentrator-5 (Zymo Research) following the protocol for plasmid cleanup, and eluted into 12 μL of elution buffer. Cleaned products were quantified using Qubit dsDNA HS Assay Kit (ThermoFisher Scientific).

Each assembled, cleaned helper plasmid mixture was then transformed into NEB Stable Competent *E. coli* (New England Biolabs) using 25 μL heat-shock reactions according to the manufacturer’s protocol. The recovery culture (1 mL) for each tile was used to inoculate a 30 mL liquid broth culture containing LB and 50 μg/mL kanamycin, which was incubated overnight (16-18 h) at 37°C with shaking. A sample of the recovery was used for serial dilutions and plated on LB-kanamycin plates that incubated overnight (16-18 h) at 37°C. Dilution plates were counted to estimate the number of transformants and coverage of variants in each tile’s culture. A total of 6 mL from each liquid culture was used to extract the plasmid from bacterial cells using the GeneJET Plasmid Miniprep kit (Thermo Scientific), according to manufacturer’s instructions. Prepared plasmids were quantified using Qubit dsDNA BR Assay Kit (ThermoFisher Scientific). Whole plasmid sequencing was performed by Plasmidsaurus using Oxford Nanopore Technology with custom analysis and annotation.

#### Transfer of assembled ORFs from helper vector into assay vectors

For each gene, all assembled helper vectors were pooled in equimolar amounts for a final transfer into barcoded assay vectors via Golden Gate assembly with SapI (New England Biolabs). For genes in which only a portion of the ORF was assembled in the helper vector, additional WT gene blocks comprising the sequences flanking the variant region, cloned in a different helper vector (pTwist Kan High copy v2), were included to complete the full-length ORF. Each reaction consisted of: 5 μL 10X rCutSmart Buffer, 5 μL ATP (10 mM) (New England Biolabs), 2.5 μL SapI, 0.5 μL T4 DNA ligase, 250 ng barcoded plasmid, an appropriate amount of insert DNA (pooled assembled helper plasmid and 0-2 gene blocks at a 2:1 insert:vector molar ratio), and nuclease-free water to bring the volume to 50 μL total. Reactions were run using the following program: 15 cycles of 37°C for 5 min followed by 16°C for 5 min, then a heat-inactivation at 65°C for 15 min and 80°C for 15 min, and a hold at 4°C. While the program ran, 1 μL of SapI was added to each reaction every h during cycling, for a total of 2 additions. When the program was complete, products underwent an additional digest with SapI and Exonuclease V. The following was added directly to each reaction: 1 μL SapI, 0.5 μL Exonuclease V, 0.5 μL 10X rCutSmart Buffer, and 2.5 μL nuclease-free water. The additional digest was incubated at 37°C for 2 h, followed by heat inactivation at 80°C for 10 min, and a hold at 4°C. Reactions were cleaned using Zymo DNA Clean & Concentrator-5 (Zymo Research) following the protocol for plasmid cleanup, and eluted using 12 μL of elution buffer. Cleaned products were quantified using Qubit dsDNA HS Assay Kit.

Assembled, cleaned library plasmids were then used to transform NEB 10-beta Electrocompetent *E. coli* (New England Biolabs) in 25 μL reactions according to the manufacturer’s protocol using a BioRad Gene-Pulser electroporator set at 2.0kV. The recovery culture (1 mL) was bottlenecked by using various recovery volumes (500 μL, 250 μL, 125 μL, etc.) to inoculate separate 250 mL liquid cultures containing LB and 100 μg/mL carbenicillin (Teknova). A 1:1000 dilution of these cultures was plated on LB-ampicillin plates as a proxy for the number of unique molecules transformed and to estimate the barcode coverage of the library. Plates were grown at 37°C overnight, and liquid cultures were grown at 30°C overnight (16-18 h) with shaking. After overnight incubation, dilution plates were counted to estimate the coverage of barcodes in the culture; liquid cultures which had coverage between 25-50X barcodes per variant were centrifuged at 4,000 x *g* for 20 min at 4°C and plasmid DNA was isolated using the ZymoPURE II Plasmid Maxiprep Kit (Zymo Research) according to manufacturer’s instructions. Library plasmids were quantified using Qubit dsDNA BR Assay Kit. In the case where two or more library bottlenecks fell within the desired coverage of barcodes, all bottleneck preparations moved on to sequence confirmation and barcode-variant coverage estimation via sequencing as described in the next section.

#### Barcode-variant coverage estimation

Barcoded library plasmids were prepared for sequencing to confirm barcode-to-variant coverage previously estimated by colony counts. The barcode region for each plasmid was amplified by PCR, first using a unique combination of custom indexing primers (JJS16 and JJS9, **Supplementary Table 4**) at a final concentration of 0.5 μM each with plasmid DNA, 25 μL Q5 Hot Start HIgh Fidelity polymerase (New England Biolabs), and nuclease-free water to bring the volume to 50 μL. Cycling conditions were an initial denaturing at 98°C for 30 seconds, followed by 5 cycles of 98°C for 30 seconds, 64°C for 10 seconds, and 72°C for 30 seconds, followed by a final extension for 2 min at 72°C and storage at 4°C. PCR products were then cleaned using 1.5X SeraMag beads (Cytiva) and eluted in 17 μL nuclease-free water, following the manufacturer’s protocol.

For the second amplification via real-time qPCR, 2 μL of the eluted PCR product was combined with 25 μL Q5 Hot Start High Fidelity 2X Master Mix, 0.3 μL SYBR green I (ThermoFisher Scientific), primers JJS16 and JJS24 (final concentration 0.5 μM each), and nuclease-free water to bring the total volume to 50 μL. Samples were run on a CFX Connect (Bio-Rad) for a maximum of 15 cycles or until all samples reached 3,000 relative fluorescence units (RFU). Reactions were denatured at 98°C for 30 seconds, and cycled at 98°C for 30 seconds, 63°C for 10 seconds, and 72°C for 30 seconds. Products were cleaned using 1.5X SeraMag beads (Cytiva) and eluted in 17 μL nuclease-free water, then quantified using the Qubit dsDNA HS Assay Kit (ThermoFisher Scientific), pooled in equimolar concentrations, and sequenced on an Illumina NextSeq 2000 instrument using a NextSeq 1000/2000 P3 50 Cycle kit (Illumina) and using custom primers (JJS25-28). Sequencing reads were converted to FASTQ format and demultiplexed using bcl2fastq (v2.20), forward and reverse barcode reads were paired using PEAR (v0.9.11), and unique barcodes were counted^97^. Counts per barcode were plotted as a histogram and the number of barcodes in the library was estimated to confirm that bottlenecking achieved approximately 30 barcodes per variant.

### Barcode-Variant Mapping

#### Long-read sequencing

Library plasmids were digested with NheI-HF (New England Biolabs) in a reaction containing 500 ng of plasmid, 5 μL 10X rCutSmart Buffer (New England Biolabs), and nuclease-free water to bring the volume to 50 μL. The digest was prepared for long-read sequencing using the PacBio SMRTbell Prep Kit 3.0 (PacBio) with SMRTbell adapter indices. Indexed plasmids were pooled according to plasmid size and target number of variants in the library to ensure appropriate coverage via long-read sequencing. Indexed and pooled samples were then submitted to the University of Washington Long Read Sequencing Center, where they were run on a PacBio Revio instrument using a SMRT Cell 25M with a 30 h movie time and 1.417 h pre-extension time.

#### Mapping

The resulting fastq.gz files were analyzed with Pacybara^98^. The clusters_transl_softfilter.csv output file for each library was used as input for downstream LABEL-seq analyses. This file only contains barcodes which are associated with a single variant in at least 67% of reads. The PacBio fastq files and maps were also compared to the programmed oligo library to evaluate the efficacy of the library cloning methods (**Extended Data Fig. 2**).

### Cell Culture

#### Generating HaloTag RPLP

A clonal cell line stably expressing a tdMCP-tagged barcode competitor (Myc-tdMCP-HaloTag) was generated by lentiviral transduction of unrecombined HEK293 RPLP cells^7^ with a transgene cassette encoding Myc-tdMCP-HaloTag, the gene required for Zeocin (bleomycin) resistance, and an mTagBFP2 fluorescent reporter (**Supplementary Table 5**). Three days post-transduction, single cells were seeded into 96-well plates by limiting dilution in growth media supplemented with doxycycline (DOX, 2 μg/mL), Zeocin (100 μg/mL), and hygromycin B (200 μg/mL). The resultant clones were expanded and screened by flow cytometry. Clonal lines exhibiting >99% mTagBFP2 positivity were considered for downstream use. A single validated clone (clone 3) was selected for further expansion and used exclusively in all subsequent abundance assays.

#### Generating CIAR RPLP

A clonal cell line stably expressing a chemically-inducible activator of Ras (CIAR) was generated by lentiviral transduction of unrecombined HEK293 RPLP cells^7^ with a transgene cassette encoding CIAR-4a^94^, the gene required for Zeocin (bleomycin) resistance, and an mTagBFP2 fluorescent reporter (**Supplementary Table 5**). Three days post-transduction, growth media was supplemented with doxycycline (DOX, 2 μg/mL), Zeocin (100 μg/mL), and hygromycin B (200 μg/mL). Two weeks post-transduction, single cells were seeded into 96-well plates by limiting dilution. The resultant clones were expanded and screened by flow cytometry. Clonal lines exhibiting >99% mTagBFP2 positivity were considered for downstream use. A single validated clone was selected for further expansion and used exclusively in all subsequent abundance assays.

#### Generating LZTR1 knock out line

To profile KRAS variant abundance in the absence of LZTR1, a clonal LZTR1 knock out (LZTR1ko) line was generated using unrecombined HaloTag RPLP cells and a previously reported guide RNA (sgLZTR1.620)^99^. Approximately 1×10^6^ HaloTag RPLP cells were transfected with 1.5 μg of PX459-sgLZTR1.620 encoding guide, SpCas9, and a puromycin selectable marker (**Supplementary Table 5**), 0.2 μg pMAX-GFP, 7.125 μL Fugene HD and 100 μL serum-free DMEM. 24 h after transfection, media was supplemented with 2 μg/mL puromycin. Three days post-transfection, single cells were seeded into 96-well plates by limiting dilution in growth media supplemented with DOX (2 μg/mL), 200 μg/mL hygromycin, 100 μg/mL zeocin, and 100 nM SCH772984. The resultant clones were expanded and screened by flow cytometry to confirm maintenance of >98% mTagBFP2 (reporting Myc-tdMCP-HaloTag expression) and >98% EGFP (reporting presence of landing pad). Candidate clones were screened for LZTR1ko by TIDE analysis and confirmed by western blotting^100^.

#### Generating library cell lines

For pathway activity profiling, stable cell lines expressing untagged variants and MS2-circRNA barcodes were generated in the HEK293 RPLP^7^. For interaction profiling, A3-3xFLAG-tdMCP-tagged variants were also expressed in the RPLP. For abundance profiling, 3xFLAG-tdMCP-tagged variants were expressed in the HaloTag RPLP described above. The following method applies to all cell lines.

Barcoded libraries were stably recombined into landing pad cell lines and maintained in high glucose DMEM supplemented with 10% FBS and 2 μg/mL DOX (0.01% DMSO final concentration). The day before transfection, landing pad cells were plated into two 15 cm dishes using dox-free media to reach ∼85% confluence at the time of transfection. On the day of transfection, the media was further supplemented with SCH772984 to a final concentration of 100 nM. Transfection mixes were prepared by combining per 15 cm dish: 16.5 μg attB-plasmid library + 3 μg pCAG-NLS-Bxb1 + 2 μg pMAX-GFP + 2 mL serum-free DMEM + 80 μL FugeneHD, then incubated at room temperature for 15 min and added dropwise to cells. 48 h after transfection cells were expanded and maintained in media supplemented with 2 μg/mL dox and 100 nM SCH772984. 4 days after transfection cells were further expanded and a sample analyzed using flow cytometry or a Countess 3 FL (Invitrogen) to quantify recombination efficiency via an mCherry reporter. 6 days after transfection, selection was initiated by further supplementing media with 10 nM AP1903. After 6-24 h, media was replaced with fresh media excluding AP1903 but including 2 μg/mL dox and 100 nM SCH772984. 24 h after initiating selection, cells were further expanded in media supplemented with 1 μg/mL puromycin, 2 μg/mL dox and 100 nM SCH772984 until aliquots were frozen.

### Standards

To enable cross-assay comparison, cell lines expressing panels of previously characterized BRAF variants spanning the full range of BRAF variant scores^7^ were cloned into barcoded assay vectors, recombined into either the RPLP (pathway activity standards) or HaloTag RPLP (abundance standards) and spiked into each sample prior to protein enrichment. For pathway activity profiling, the standards were untagged BRAF wild type, R509Y, G469T, G258E, and K601D. For abundance profiling, the standards were 3xFLAG-tdMCP-tagged BRAF wild type, E695*, I592S, G258N, S727K, and P367N.

### Abundance profiling

#### Overview

For variant abundance assays, a tdMCP-tagged competitor protein (Myc-tdMCP-HaloTag) was co-expressed with 3xFlag-tdMCP-tagged variants and the partitioning of barcodes between the two proteins was used as a direct measurement of variant abundance. The competitor is required because barcodes are expressed at lower concentrations than some tdMCP-tagged proteins; given the high affinity of the MS2-tdMCP interaction, barcodes would otherwise be saturated with tdMCP, obscuring differences in variant abundance.

#### Cell culturing

Prior to profiling, tdMCP-tagged variant libraries were recombined into the HaloTag RPLP as described in *Generating library cell lines*. Aliquots of library and standard cell lines were thawed several days prior to each experiment and expanded to the necessary number of dishes. 3,000-5,000 cells per variant were used for each replicate. Both library and standard cell lines were thawed and maintained in high-glucose DMEM supplemented with 10% FBS, DOX (2 μg/mL) and SCH772984 (100 nM). The day after thawing, media was further supplemented with puromycin (2 μg/mL).

The day before pulldowns, cells were transferred to media containing DOX but lacking SCH772984 and puromycin. For assays with HSP90 inhibitor (HSP90i), pimitespib was added to a final concentration of 600 nM concurrently with the washout of SCH772984 and puromycin. For basal abundance assays, standard cells were added to the library culture the day prior to pulldowns. For HSP90i experiments, standard cells were added on the day of pulldowns following trypsinization but prior to centrifugation. Standard cells were added at a frequency approximately equal to that of library barcodes.

#### Lysate preparation

On the day of pulldowns, cells were collected by trypsinization and pelleted at 300 x g for 5–10 min, pooling all cells belonging to a single replicate prior to centrifugation. After pelleting, media was aspirated and cells were immediately resuspended in cold resuspension buffer (50 mM Tris pH 7.6, 150 mM NaCl, 5% glycerol, 4 mM MgCl_2_, 10 mM NaF, 1× Promega protease inhibitor, 4 μL/mL Ribolock RNase inhibitor, 300 nM dummy RNA - see reagent list in **Supplementary Table 6** for preparation) using 1 mL per 15 cm dish, with cell clumps dissociated by gentle pipetting. An equal volume of cold lysis buffer (resuspension buffer supplemented with 0.2% IGEPAL CA-630 and lacking dummy RNA) was added immediately thereafter, followed by gentle pipetting. All subsequent steps were performed on ice or at 4°C. Lysates were cleared by centrifugation at 17,000 x g for 10 min at 4°C.

#### Protein enrichment

Cleared lysate from each replicate was divided, allocating 25% to anti-FLAG (variant) enrichment and 75% to HaloLink (competitor) enrichment. Prior to use, anti-FLAG beads (supplied as a 50% slurry) and HaloLink resin (supplied as a 25% slurry) were washed 2–3 times with 10 slurry volumes of wash buffer (50 mM Tris pH 7.6, 150 mM NaCl, 5% glycerol, 4 mM MgCl_2_, 0.1% IGEPAL CA-630) then resuspended in wash buffer to their original slurry concentrations. Washed resin was added to lysate using 100 μL slurry per 1 mL lysate. Enrichment of tdMCP–RNA complexes was carried out in low-bind tubes, rotating end-over-end for 90 min at 4°C. Beads were subsequently washed three times with 10 slurry volumes of cold wash buffer without incubation. After the final wash, the buffer was removed and the beads were resuspended in 50–100 μL nuclease-free water plus 4x water volumes of TRIzol (from TRI-Isolate RNA Pure kit) then stored at −20°C until ready for further processing.

High concentrations of select protease inhibitors, including PMSF, rapidly inactivate HaloLink resin and severely decrease the efficiency of competitor enrichment. To minimize this issue, we used a protease cocktail from Promega which includes inhibitor concentrations compatible with HaloLink resin.

### Pathway activity profiling

#### Overview

For pathway activity assays, a reporter protein (3xFLAG-tdMCP-ERK2 K54R) bound to variant-specific barcodes was selectively captured using DARPin reagents that distinguish phosphorylated ERK (pERK) from unphosphorylated ERK (uERK); the ratio of pERK to uERK barcodes provides the pathway activity score for each variant. All assays included internal activity standards.

#### Generating pathway activity affinity reagents

Plasmids encoding His- and AviTagged-E40 and pEM1 DARPins^101^ were transformed into *E. coli* BL21(DE3) and grown in LB–ampicillin at 37°C. Protein expression was induced at OD₆₀₀ ∼ 0.6 with 0.5 mM IPTG and grown for 4-5 h in the presence of 12.2 mg/L biotin. Cells were collected and lysed in TBS supplemented with 5 mM imidazole, 5% glycerol, 2 mM β-mercaptoethanol, and a Pierce protease inhibitor tablet. His-tagged protein was purified by Ni–NTA affinity chromatography with sequential high-salt and imidazole washes, eluting in 250 mM imidazole. Purified DARPins were dialyzed into storage buffer (25 mM Tris pH 8.0, 150 mM NaCl, 5% glycerol), quantified by Coomassie against a BSA standard, and stored as single-use aliquots at −80°C.

#### Cell culturing

Prior to profiling, untagged variant libraries were recombined into the HEK293 RPLP as described in *Generating library cell lines*. Aliquots of library and standard cell lines were thawed and expanded several days prior to each experiment and maintained in high-glucose DMEM supplemented with 10% FBS, DOX (2 μg/mL) and SCH772984 (100 nM). The day after thawing, media was further supplemented with puromycin (2 μg/mL).

Three days before pulldowns, library cells were seeded into three 15 cm dishes at 50-80% confluency. Standard cells were added to these cultures at a frequency approximately equal to that of library barcodes. Each plate was transduced using frozen aliquots of lentivirus harboring the pErk reporter transgene, 3xFLAG-tdMCP-ERK2 K54R. Beginning two days before pulldowns, cultures were expanded or passaged 1:2. The day before pulldowns, cells were expanded and transferred to media containing DOX but lacking SCH772984 and puromycin. Plates were seeded at a confluence that achieved 1,800-4,000 cells per variant per replicate on the day of pulldowns. Samples were also analyzed by flow cytometry to confirm >60% transduction efficiency.

#### DARPin bead preparation

The day prior to pulldowns, pERK (pEM1) and uERK (E40) DARPin beads were prepared. Biotinylated pEM1 and E40 DARPins were conjugated to streptavidin agarose by diluting each DARPin in TBS (pH 7.6) to a final concentration of 32 μM in 1 mL total volume, then adding 300 μL high-capacity streptavidin agarose (supplied as a 50% slurry). Conjugation was carried out by end-over-end rotation at 4°C overnight in a low-bind Eppendorf tube. On the day of pulldowns, DARPin-conjugated beads were centrifuged at 300 x g for 1 min, washed once with 10 slurry volumes of TBS to remove unbound DARPin, centrifuged again, then resuspended in 150 μL TBS yielding 300 μL of 20× concentrated DARPin-conjugated resin. This volume of resin was used to enrich phosphorylated and unphosphorylated ERK from 6 mL of lysate (three 15 cm dishes) and was scaled proportionally for larger experiments.

#### Lysate preparation

On the day of pulldowns, cells were collected by trypsinization and pelleted at 300 x g for 5–10 min, pooling all cells belonging to a single replicate prior to centrifugation. After pelleting, media was aspirated and cells were immediately resuspended in cold resuspension buffer (50 mM Tris pH 7.6, 150 mM NaCl, 5% glycerol, 4 mM MgCl_2_, 10 mM NaF, 1× Pierce protease inhibitor, 2 mM PMSF, 4 μL/mL Ribolock RNase inhibitor, 300 nM dummy RNA - see reagent list in **Supplementary Table 6** for preparation, 100 μL/10 mL buffer phosphatase inhibitor cocktails 2 and 3) using 1 mL per 15 cm dish, with cell clumps dissociated by gentle pipetting. An equal volume of cold lysis buffer (resuspension buffer supplemented with 0.2% IGEPAL CA-630 and lacking dummy RNA) was added immediately thereafter, followed by gentle pipetting. All subsequent steps were performed on ice or at 4°C. Lysates were cleared by centrifugation at 17,000 x g for 10 min at 4°C.

#### Protein enrichment

Two sequential protein enrichments were subsequently performed. First, unphosphorylated tdMCP-ERK2 was enriched by incubating cleared lysates with the E40-conjugated streptavidin beads, using 50 μL prepared bead slurry per 1 mL lysate, equivalent to a final concentration of 4 μM E40. This enrichment was performed in low-bind Eppendorf tubes, rotating end-over-end for 1 h at 4°C. After enrichment, beads were separated from lysate by centrifugation at 300 x g for 1 min at 4°C. The supernatant was transferred to a fresh low-bind Eppendorf tube and the beads reserved on ice. To enrich the phosphorylated fraction of tdMCP-ERK2, pEM1-conjugated streptavidin beads were added to the supernatant using 50 μL prepared bead slurry per 1 mL lysate, equivalent to a final concentration of 0.5 μM pEM1. These mixtures were rotated end-over-end for 1 h at 4°C. During the pEM1 incubation, E40 beads were washed 3 times using a 10x bead slurry volume of cold wash buffer (50 mM Tris pH 7.6, 150 mM NaCl, 5% glycerol, 4 mM MgCl_2_, 0.1% IGEPAL CA-630) without incubation. After each wash, beads were separated from the wash buffer by gentle centrifugation, 300 x g for 1 min. After the final wash, the buffer was removed and the beads were resuspended in 50–100 μL nuclease-free water plus 4x water volumes of TRIzol (from TRI-Isolate RNA Pure kit) then stored at −20°C until ready for further processing. pEM1 beads were similarly washed and stored.

### Interaction profiling

#### Cloning A3-KRAS library

3xFLAG–tdMCP was excised from the KRAS abundance library by digestion with AlfII and BamHI and replaced with an A3–3xFLAG–tdMCP insert via Gibson assembly (10 ng linearized backbone, 2 ng insert). The assembled product was transformed into NEB 10-beta electrocompetent *E. coli* following the manufacturer’s protocol and expanded overnight in LB with ampicillin. Plasmid DNA was isolated by midiprep and validated by Sanger sequencing to confirm cassette integration. The plasmid pool was sequenced using Illumina sequencing to confirm retention of KRAS barcode coverage.

#### Cell culturing

Prior to profiling, A3-3xFLAG-tdMCP-tagged variant libraries were recombined into HEK293 RPLP cells as described in *Generating library cell lines*. An aliquot of the library cell line was thawed and expanded several days prior to each experiment, maintaining in high-glucose DMEM supplemented with 10% FBS, DOX (2 μg/mL) and SCH772984 (100 nM). The day after thawing, media was further supplemented with puromycin (1 μg/mL).

Lentivirus encoding BirA–LZTR1 was produced in HEK293T cells using a third-generation system by transfecting 15 cm dishes with 10 µg transfer plasmid, 9 µg gag/pol, 4.5 µg Rev, and 4.5 µg VSV-G at a 1:3 DNA:PEI mass ratio. Virus-containing media was collected 48 h later, clarified, 0.45 µm–filtered and immediately used.

Three days before pulldowns, one 15 cm dish of the A3-KRAS library (85% confluent) was transduced using 12 mL filtered viral media. To minimize background labeling, 24 h after transduction cells were transferred to media prepared using dialyzed FBS. 48 h after transduction, cells were transferred to media containing DOX but lacking SCH772984 and puromycin. 72 h after transduction (on the day of the pulldowns), 15 cm dishes of cells were treated with 200 µM biotin for 15 min at 37°C.

#### Lysate Preparation

To halt biotin labeling, cells were collected by trypsinization and pelleted at 300 x g for 5–10 min, pooling all cells belonging to a single replicate prior to centrifugation. After pelleting, media was aspirated and cells were immediately resuspended in cold resuspension buffer (50 mM Tris pH 7.6, 150 mM NaCl, 5% glycerol, 4 mM MgCl_2_, 10 mM NaF, 1× Pierce protease inhibitor, 2 mM PMSF, 4 μL/mL Ribolock RNase inhibitor, 300 nM dummy RNA - see reagent list in **Supplementary Table 6** for preparation) using 1 mL per 15 cm dish, with cell clumps dissociated by gentle pipetting. An equal volume of cold lysis buffer (resuspension buffer supplemented with 0.2% IGEPAL CA-630 and lacking dummy RNA) was added immediately thereafter, followed by gentle pipetting. All subsequent steps were performed on ice or at 4°C. Lysates were cleared by centrifugation at 17,000 x g for 10 min at 4°C.

#### Protein Enrichment

Cleared lysate from each replicate was divided, allocating 25% to anti-FLAG (total variant) enrichment and 75% to streptavidin (labeled variant) enrichment. Prior to use, anti-FLAG beads and streptavidin resin (both supplied as 50% slurries) were washed 2–3 times with 10 slurry volumes of wash buffer (50 mM Tris pH 7.6, 150 mM NaCl, 5% glycerol, 4 mM MgCl_2_, 0.1% IGEPAL CA-630) then resuspended in wash buffer to their original slurry concentrations. Washed resin was added to lysate using 100 μL (anti-FLAG) or 50 μL (streptavidin) slurry per 1 mL lysate. Enrichment of tdMCP–RNA complexes was carried out in low-bind tubes, rotating end-over-end for 60 min at 4°C. Beads were subsequently washed three times with 10 slurry volumes of cold wash buffer (50 mM Tris pH 7.6, 150 mM NaCl, 5% glycerol, 4 mM MgCl_2_, 0.1% IGEPAL CA-630) without incubation. After the final wash, the buffer was removed and the beads were resuspended in 50–100 μL nuclease-free water plus 4x water volumes of TRIzol (from TRI-Isolate RNA Pure kit) then stored at −20°C until ready for further processing.

### Pathway activity and abundance profiling after SOS1 activation

To profile pathway activity or abundance under SOS1 activation, an engineered chemically-induced activator of Ras (CIAR) switch, NS3a-CIAR, was used^94^. In this system, the constitutively active RAS-activating module of SOS1, comprising the REM and Cdc25 domains, are flanked by NS3a and the NS3a-binding peptide ANR which, together, occlude the SOS1 active site. Addition of small-molecule inhibitors of NS3a, such as danoprevir, displaces the ANR peptide allowing SOS1 to activate cellular Ras.

For abundance profiling under SOS1 activation, variant libraries in the HaloTag RPLP were plated into 15 cm dishes three days prior to pulldowns. Two days prior to pulldowns, each 15 cm dish was transfected with 18 μg CIAR 4a (**Supplementary Table 5**), 2 μg pMAX-GFP, 75 μL Fugene HD, and 1800 μL serum-free DMEM. One day prior to pulldown, the growth media was replaced with media containing DOX (2 μg/mL) and 1 μM danoprevir. Variant profiling was subsequently performed as described in *abundance profiling*.

For pathway activity profiling under SOS1 activation, untagged variant libraries were recombined into the CIAR RPLP cell line. The day prior to pulldown, the growth media was replaced with media containing DOX (2 μg/mL) and 1 μM danoprevir. Variant profiling was subsequently performed as described in *pathway activity profiling*.

### Amplicon generation and sequencing

#### RNA extraction and cleanup

To prepare barcodes for sequencing, RNA was extracted from the affinity reagent usingTRI-Isolate RNA Pure Kit and eluted into 53 μL nuclease-free water. RNA was further concentrated using Zymo RNA Clean & Concentrator columns, eluted into 10-25 μL nuclease-free water, and stored at −80°C. Prior to reverse transcription, RNA was quantified using the Qubit^TM^ RNA High Sensitivity assay kit.

#### Reverse transcription

Following the manufacturer’s instructions, concentrated and quantified RNA was reverse transcribed using SuperScript™ IV Reverse Transcriptase and custom primer JJS1. All RNA from a given enrichment was used in the reaction. As an example, the following volumes are appropriate for 100 ng of input RNA and were scaled according to the specific experiment. Primer/template mixes corresponding to a single enrichment were generated by combining concentrated RNA + 0.75 μL JJS1 (2 μM) + 0.75 μL dNTP’s (10mM ea.) + nuclease-free water to a final volume of 9.75 μL. RNA denaturation and primer annealing was performed by heating mixes to 75°C for 5 min then setting on ice. 0.75 μL DTT (0.1M) + 0.75 μL Ribolock RNase inhibitor + 0.75 μL SSIV enzyme + 3 μL SSIV buffer was subsequently added to each reaction (total volume = 15 μL). cDNA was generated by heating the reaction at 57°C for 30 min, 80°C for 10 min, then holding at 4°C. Reverse transcription reactions were stored at −20°C until ready to use.

#### PCR

cDNA was cleaned using Zymo DNA Clean & Concentrator columns (5 μg or 25 μg capacity columns) and eluted into 45 μL nuclease-free water. cDNA was either stored at −20°C or immediately used in PCR round #1 to generate double stranded DNA and add amplicon adapters, amplifying only cDNA derived from circRNA. Specifically, cDNA was split into two PCR reactions, each including 20 μL cleaned cDNA + 2.5 μL JJS8 (10 μM) + 2.5 μL JJS9 (10 μM) + 1.5 μL DMSO + 25 μL Q5® Hot Start High-Fidelity 2X Master Mix. Cycling included 98°C for 30 s, five cycles of (98°C for 10 s > 64°C for 20 s > 72°C for 30 s), 72°C for 2 min, and 4°C hold. Replicate reactions were pooled then cleaned using Zymo DNA Clean & Concentrator columns (5 μg capacity) and eluted using 25 μL nuclease-free water. Illumina adapters were added in a second round of PCR. This reaction included 5-20 μL of cleaned reaction #1 + 2.5 μL JJS16 (10 μM) + 2.5 μL JJS24 (10 μM) + 0.5 μL 10x Sybr gold + 25 μL Q5® Hot Start High-Fidelity 2X Master Mix + 1.0 μL DMSO + nuclease-free water to 51.5uL. PCR cycling included one round at 98°C for 30s, then cycling at (98°C for 10 s > 63°C for 20 s > 72°C for 30 s). Reaction progress was monitored on a BioRad CFX Connect and stopped when RFU reached ∼3,000 (∼10-15 cycles). Amplicons were pooled based on final RFU values, separated by gel electrophoresis and the band at ∼220 bp excised using Freeze ‘N Squeeze™ DNA Gel Extraction Spin Columns. The DNA was further concentrated using Zymo DNA Clean & Concentrator columns and quantified using Qubit™ dsDNA HS Assay Kit. Amplicons were sequenced as described in *Illumina Sequencing*.

#### Illumina sequencing

Amplicons were sequenced on a NextSeq 2000 using NextSeq 1000/2000 P3 Reagents. 16 nucleotide barcodes were read twice by the custom paired-end sequencing primers JJS25 and JJS26 and 8 nucleotide dual indices were sequenced using JJS27 and JJS28.

### Data processing and variant scoring

Sequencing reads were converted from BCL to FASTQ format and demultiplexed using bcl2fastq (v2.20). Paired sequencing reads were subsequently joined using PEAR (v0.9.11) and the assembly parameters -q 30 -j 8 -n 10. The resulting paired-end FASTQ files were then processed as follows. Raw sequencing count data for each barcode was queried against the lookup table of barcode-variant pairs (see *Mapping* section) to retrieve the corresponding variant. Barcodes mapping ambiguously to a variant and standard were excluded from all downstream analyses.

#### Barcode filtering

For each assay type, sequencing counts derived from two pulldowns were used to compute a per-barcode enrichment ratio. For pathway activity, the ratio was defined as pEM1 counts divided by E40 counts; for protein abundance assays, as anti-FLAG counts divided by anti-HaloTag counts; and for protein–protein interaction assays, as anti-strep counts divided by anti-FLAG counts. A minimum read count threshold of 10 was applied to both the numerator and denominator; barcodes with counts below this threshold in either channel were excluded from ratio calculation.

To remove barcodes exhibiting anomalously high or low enrichment ratios inconsistent with the behaviour of other barcodes representing the same variant (potentially reflecting RT-PCR or enrichment artifacts), a biased barcode filtering step was performed. For each variant and replicate, the per-barcode ratio was compared against the median ratio of all barcodes for that variant; barcodes with a ratio 2.5 standard deviations above the variant median were flagged. Barcodes flagged in two or more replicates were excluded from all replicates. Following filtering, per-barcode enrichment ratios were recalculated. Each barcode’s enrichment ratio was then normalized to the mean WT enrichment ratio within the same replicate to yield a per-barcode functional score. Only barcodes linked to WT, synonymous, missense, 3 nucleotide deletion variants (which are referred to as deletions here), and nonsense variants that were programmed in the library generation step were retained for further analysis.

#### WT-relative scoring

Barcode-level scores were aggregated to the variant level by computing the mean score across all retained barcodes for each variant within each replicate. A variant-level average score was then calculated as the mean of the three replicate scores. Variants were retained for downstream analysis only if they were supported by a mean of at least 5 quantified barcodes across the three replicates.

Two of 192 replicates (MRAS activity rep 2, KSR1 C-terminal activity rep 3) had compressed variant effect scores relative to the other two replicates of the same library and assay, with scores about half as far from WT on a log-scale. We modeled each replicate’s log-transformed scores as the product of a scaling factor and the variant’s true effect, plus noise. We estimated each replicate’s scaling factor from the covariances of log-transformed scores between each pair of replicates. We adjusted the two compressed replicates using their scaling factors (1.98 for MRAS activity, 2.00 for KSR1 C-terminal activity). This procedure does not change the WT score. Log-transformed scores below the 1st and above the 99th percentile were instead scaled by a fixed factor equal to the fold-change at that percentile. All analyses presented here use these adjusted replicate scores; **Supplementary Table 2** reports both adjusted and unadjusted replicate scores.

#### Standardized scoring

To enable quantitative comparison of functional scores across libraries, assays, and experimental batches, per-replicate scores were calibrated against a panel of spiked-in standard variants with pre-assigned functional scores. For pathway activity assays, standards spanning the dynamic range of the assay were included: BRAF R509Y (score = 0.40), BRAF WT (score = 1.00), BRAF G469T (score = 8.36), BRAF G258E (score = 12.0), and BRAF K601D (score = 18.0). For protein abundance assays, standards were: BRAF E695* (score = 0.16), BRAF I592S (score = 0.29), BRAF G258N (score = 0.53), BRAF S727K (score = 0.66), BRAF P367N (score = 0.71), and BRAF WT (score = 1.00). For each replicate, a linear regression with the intercept constrained to zero was fit between the measured scores and the assigned standard scores; the resulting slope was used to transform all variant scores into standardized units. The per-replicate standardized scores were then averaged to yield a final standard-adjusted score for each variant. Variants were retained for downstream analysis only if they were supported by a mean of at least 5 quantified barcodes across the three replicates.

#### Variant classification and structural annotation

To classify variants by functional consequence, score thresholds were derived empirically from the distribution of synonymous WT variant scores within each experiment. Variants with scores below the 2.5th percentile or above the 97.5th percentile of the synonymous WT distribution were classified as functionally altered (reduced or increased activity or abundance), while those within these bounds were classified as WT-like.

Positions with DSSP v3.1.4-computed relative solvent accessible surface area (rSASA) below 0.25 were considered buried^102^. Interdomain contacts were defined as positions with any heavy atom within 4 Å of a heavy atom in a different curated domain of the same protein. Where categories overlap, interdomain contact takes precedence over burial.

### Dominant negative analysis

For each activity assay, pathway activity scores for barcodes corresponding to non-protein-expressing (empty) vectors were computed and then used to generate 500 empty vector pathway activity scores by averaging 10 randomly selected barcodes (with replacement) at a time. The 2.5th percentile of these 500 scores was computed and this procedure was repeated with 200 random seeds. Variants whose pathway activity fell below the median of these 200 2.5th percentile values were classified as dominant negative in each library and treatment. Because overexpression of WT GRB2, KSR1, KSR2, MEK1 and MEK2 led to lower pathway activity than the empty vector baseline, we did not attempt to identify dominant negative variants in these targets.

Missense and single-position deletion dominant negative variants were assigned to three structural categories: active site, buried non-active site, or interface. A variant was classified as residing in an active site if it was at a position carrying an active-site feature, curated by hand from NCBI Conserved Domain Database annotations; buried non-active site if it was at a non-active-site position with relative solvent-accessible surface area (rSASA) below 0.25; and interface if it was at a position within 4 Å heavy atom distance of a chain of an interacting protein in a curated set of PDB complexes. DSSP v3.1.4 was run on full-length monomer models from the AlphaFold Database (release v4, AlphaFold Monomer v2.0 pipeline) to generate both secondary structure and rSASA annotations.

Amino acid positions were mapped to GRCh38 coordinates and phyloP^103^ 100-way vertebrate scores were retrieved from the UCSC REST API. The position-level phyloP score is the mean phyloP over the bases of the codon encoding that position. Kinase conservation scores were computed on the human kinome alignment^90^ by use of a previously described method^91^.

Scored single nucleotide missense and nonsense variants were tested for whether they had been observed in gnomAD v4 (accessed 1 April 2026), All of Us Curated Data Repository v7 (accessed 11 July 2025) and AACR GENIE v19.0 via cBioPortal (accessed 8 April 2026)^56,59,104^. A variant counted as observed at an allele count above zero in gnomAD and All of Us, or in two or more tumours in GENIE. Observation rates were compared between pathway-activity classes by two-sided Fisher’s exact test with WT-like variants as the reference class, using conditional maximum-likelihood odds ratios with exact 95% confidence intervals and Benjamini–Hochberg correction.

### HSP90-inhibited abundance score analysis

Standard variants were used to place abundance measurements on a common internal-standard scale, giving standardised abundance scores for 105,974 variants in both pimitespib (HSP90 inhibitor, HSP90i)-treated and inhibitor-free, basal conditions. Standardised abundance scores in HSP90i and basal conditions were compared across missense, deletion, and nonsense variants for each protein.

Variants were classified as reduced-, increased-, or WT-like abundance in each condition using the synonymous distribution of the same target and condition (<2.5th percentile of synonymous scores = reduced, >97.5th percentile = increased, 2.5th-97.5th percentile = WT-like). Separately, a variant was classified as dependent on HSP90 if the difference between its standardized abundance in the control and HSP90i conditions was at least as large as the difference between the WT score and the reduced-abundance threshold (i.e. 2.5th percentile of synonymous scores). Variants were also classified as buffered or poorly buffered. A variant was classified as buffered if it had WT-like or increased abundance in the basal condition and reduced abundance in the HSP90i condition; poorly buffered if it had reduced abundance in both conditions; and WT-like/increased if it did not have reduced abundance in either condition. 3.2% of missense variants had reduced basal abundance but WT-like or increased HSP90i abundance; these variants were excluded from the analyses presented in **Figure 5** and **Extended Data Figure 8**.

Kinase-domain boundaries follow the curated per-protein domain annotation described in the *Dominant negative analysis* section. Using the human kinome alignment^90^, per-position fractions of each buffering class were averaged across the nine profiled kinases by alignment column and projected onto the Aurora-A kinase structure 3E5A^105^. Kinase conservation and phyloP were computed as above. ΔΔG values were predicted with SPURS^69^, and rSASA annotations were generated as above.

A broader set of candidate features of buffering was assembled for kinase-domain missense variants, spanning substitution chemistry, burial and packing, secondary structure and kinase lobe, kinase-motif membership, geometry, interfaces and chaperone contacts. Substitution chemistry features were gathered from amino acid properties and physicochemical (changes in hydrophobicity^106^, changes in side chain volume^107^, changes in polarity^108^) and substitution-matrix tables (BLOSUM62^109^, Grantham distance^110^). Burial and packing terms, like rSASA, were computed using DSSP v3.1.4 on AlphaFold Database v4 monomer models. Secondary structure, kinase lobe, kinase-motif membership and distances to the catalytic-cleft motifs were assigned from the human kinome alignment^90^ together with the structural models. As above, interface positions were defined as those within 4 Å heavy atom distance of a chain from an interacting protein in a curated PDB complex. Chaperone-contacting positions were defined as interface positions in BRAF or CRAF complexes with HSP90 and CDC37 (PDB 7ZR0 and PDB 8U1L)^111,112^, and transferred to the remaining kinases using the human kinome alignment^90^. HSP90 contacts were divided into contacts on the unfolded N-lobe and contacts on the remaining folded portion.

For each target-feature combination, a univariate multinomial logistic regression was fitted using kinase-domain missense variants to predict buffering class with WT-like/increased as the reference class. Continuous predictors were expressed as z-scores computed over variants from all targets so that coefficients represent the change in log-odds per standard deviation and are comparable across targets. The logistic regression coefficient for buffered versus WT-like/increased variants for each target-feature combination is reported, ordered by decreasing mean McFadden pseudo-R².

### Interactor analysis

Complexes containing target proteins were assembled from three sources. Experimental structures were retrieved from the RCSB PDB, restricted to complexes solved by X-ray diffraction at ≤ 3.5Å or by cryo-electron microscopy at ≤ 4.5Å. To prevent repeated copies from entries with more than two chains within an asymmetric unit from generating redundant pairs, chains were grouped by interacting partner and one representative heterodimer was chosen for each target protein-interacting partner combination. For interacting pairs with many structures in the PDB (e.g. the BRAF dimer), the five structures with the highest resolution were retained. This yielded 343 complexes from 266 PDB entries. RoseTTAFold2-PPI predictions were obtained from a published proteome-scale screen filtered to interactions reported at an FDR of 0.2. Structures for these interactions were the indicated best AlphaFold2 or AlphaFold-Multimer models from the screen, giving 355 complexes containing scored targets^76^. Lastly, AlphaFold-Multimer predictions (comprising three ranked models for each interacting pair) were obtained from a published proteome scale screen, filtered to 771 complexes over 257 interacting pairs containing scored targets and with SPOC score ≥ 0.69 corresponding to an FDR of 0.2^113^. Overall, this yielded 1,469 complexes containing at least one scored chain.

Complexes were first processed using DSSP v3.1.4 to annotate solvent-exposed positions, defined as positions with rSASA ≥ 0.25. Interface positions were then defined as positions with less than a 6 Å inter-chain distance between Cβ (or Cα if the amino acid at that position was glycine). We used Cβ rather than heavy atom distances because structure predictors model backbone positions more accurately than side-chain conformations^114,115^. “Control” positions were defined as solvent-exposed positions within an annotated domain on the scored chain with at least 10 Å Cβ intra-chain separation from the nearest interface position. For predicted complexes with reported pLDDT, low confidence positions with pLDDT < 60 were also removed. Complexes were then filtered to those that had at least 3 scored interface and 3 scored control positions on the target and no confident inter-chain Cβ clash, defined as any high-confidence position with less than 3 Å inter-chain separation. This yielded 928 assessable complexes spanning 395 protein pairs.

For each assessable complex and scored target protein, the WT-relative activity and abundance scores of missense variants at interface positions were compared to those at control positions, yielding a two-sided Mann-Whitney U-test p-value and Cohen’s d score (a measure of effect size) for each assay. Heterocomplexes with two scored targets were tested twice, once for each target. The Benjamini-Hochberg False Discovery Rate (BH-FDR) correction was then applied to these p-values across all complexes within each assay to give q-values. Functionally supported complexes were defined as complexes with q < 0.05 and |Cohen’s d| ≥ 0.2.

For display in the volcano plots only (**Fig. 6c,d**), complexes of the same protein pair produced by the same predictor were merged into one point when their interfaces overlapped to an intersection-over-union (IoU) of at least 0.3. Each merged point takes the median Cohen’s d and the geometric mean of the p-values of its complexes. Then, the Benjamini-Hochberg correction was applied across the merged points within each assay. The results were displayed as volcano plots of q-value against Cohen’s d.

To produce **Extended Data Fig. 9i-k**, interactors with at least five scored interface positions were clustered by where they bind on each scored target. Every interacting protein, across all predictors, was assigned a footprint: the union of its interface positions on the target protein over all of its complexes, predicted or experimental. Footprints were compared by their IoU, counting a residue as shared if it lay within 6 Å Cβ distance of any residue in the other footprint, so that partners contacting the same surface patch were treated as overlapping. Footprints were clustered by complete-linkage hierarchical clustering using IoU such that cluster members shared at least 0.5 IoU (i.e. for every pair of interactors within a cluster, shared positions make up at least half of their combined footprint). Clusters were classified as singletons, containing one interacting protein, or as promiscuous, containing two or more interactors.

Interaction partners were called as reported when present in BioGRID multi-validated physical interactions (release 5.0.259)^74^ or in UniProt curated^73^ “Interacts with” annotations. These databases were also used to report the total number of reported interactions for our scored targets. Partner druggability was annotated from the Open Targets Platform^81^ using both approved drugs and clinical candidates as well as high-quality chemical probes (accessed 27 July 2026). Every molecule returned was checked by Claude Opus 5 against the primary literature to confirm it engages that partner (**Supplementary Table 3**).

